# Persistent Immune and Clotting Dysfunction Detected in Saliva and Blood Plasma after COVID-19

**DOI:** 10.1101/2022.03.18.484814

**Authors:** Hyesun Jang, Saibyasachi Choudhury, Yanbao Yu, Benjamin L. Sievers, Terri Gelbart, Harinder Singh, Stephen A. Rawlings, Amy Proal, Gene S. Tan, Davey Smith, Marcelo Freire

**Affiliations:** Genomic Medicine and Infectious Diseases, J. Craig Venter Institute, La Jolla, CA, and Rockville, MD, USA; DGG-Genomics Division, Agilent, Technologies, Inc., La Jolla, CA 92037; Department of Chemistry & Biochemistry, University of Delaware, Newark, DE, USA, 19716; MMP Adult Infectious Disease, Maine Medical Center, South Portland, ME, 04106; Division of Infectious Diseases and Global Public Health Department of Medicine, University of California San Diego, La Jolla, CA, USA; PolyBio Research Foundation. Mercer Island, WA, USA

## Abstract

A growing number of studies indicate that coronavirus disease 2019 (COVID-19) is associated with inflammatory sequelae, but molecular signatures governing the normal vs. pathologic convalescence process have not been well-delineated. We characterized global immune and proteome responses in matched plasma and saliva samples obtained from COVID-19 patients collected between 4-6 weeks after initial clinical symptoms resolved. Convalescent subjects showed robust IgA and IgG responses and positive antibody correlations between matched saliva and plasma samples. However, global shotgun proteomics revealed persistent inflammatory patterns in convalescent samples including dysfunction of salivary innate immune cells and clotting factors in plasma (e.g., fibrinogen and antithrombin), with positive correlations to acute COVID-19 disease severity. Saliva samples were characterized by higher concentrations of IgA, and proteomics showed altered pathways that correlated positively with IgA levels. Our study positions saliva as a viable fluid to monitor immunity beyond plasma to document COVID-19 immune, inflammatory, and coagulation-related sequelae.

## Introduction

Patients infected with the SARS-CoV-2 virus driving the COVID-19 pandemic generally experience a course of acute illness that lasts for approximately 2 weeks. For example, Byrne *et al.* reported that the estimated mean time from COVID-19 symptom onset to two negative PCR tests was 13.4 days (*1*). After this acute phase of illness, COVID-19 patients who do not experience further complications generally produce SARS-CoV-2 associated antibodies and enter the recovery or convalescent stage of the disease. However, despite SARS-CoV-2 antibody production and a decrease in clinical symptoms, it is possible that convalescent COVID-19 patients (3-6 weeks after initial illness) still experience immune or coagulation-related sequelae. Indeed, an increasing number of complications are being reported in individuals after acute COVID-19 (*2, 3*). One longitudinal study followed COVID-19 survivors for up to 6-, and 12-month after symptom onset. While the majority of subjects returned to normal life and produced antibody levels, they exhibited a dynamic range of recovery levels (*4*) and the complete molecular fingerprint caused by virus exposure remains unknown. It is consequently important to document molecular signatures in convalescent COVID-19 subjects to better define the normal vs. pathologic convalescence process and to detect the possible initiation of aberrant innate immune activation, especially in fluids that are in direct contact with SARS-CoV-2 (*5*).

We characterized global immune and proteome responses after SARS-CoV-2 infection in matched plasma and saliva obtained from convalescent COVID-19 subjects (n=34), with samples obtained from healthy individuals pre-COVID-19 era serving as healthy controls (n=13). We focused on the analysis of saliva in addition to plasma for the following reasons: 1) saliva is a practical and optimal body fluid to monitor for host and immune-inflammatory markers 2) saliva is a direct surrogate for SARS-CoV-2 antibody responses derived from bronchial-alveolar lymphoid tissues (BALT) (*6*), 3) saliva can reflect systemic reactions to infection since more than 90% of body protein components are detected from saliva (*7–9*), 4) saliva contains oral microbiome commensals which shape the host immune profile (*10, 11*), and 5) oral inflammation can influence the severity of systemic inflammatory responses (*12–14*). A human salivary proteome database was initially developed to explore saliva as a source of mapping markers in health and disease and to further advance comparisons to other body fluids (*15*). The current effort is now publicly available to share studies related to the composition and function of saliva biofluid (salivaryproteome.org). To date, evidence is lacking to understand immune responses (present in fluids such as saliva and plasma) on a global scale to evaluate subjects that experienced and resolved from SARS-CoV-2 infection.

Here, we measured and compared SARS-CoV-2 antibody responses in matched saliva and plasma samples, with SARS-CoV-2 S bearing pseudovirus particles used to evaluate virus neutralization. We used shotgun proteomics to analyze early immune and host cell-mediated markers that may activate inflammation and endothelial damage during COVID-19 convalescence. Last, we characterized host functional pathways impaired by SARS-CoV-2 infection by comparing responses in convalescent COVID-19 subjects to that of healthy controls and by comparing responses in saliva to that of plasma.

## Results

### The salivary antibody repertoire towards SARS-CoV-2 antigens significantly correlated with matched plasma serology

As body fluids are exposed to different antigens, we investigated how SARS-CoV-2 antibody responses in saliva compared to those in matched plasma. Paired saliva and plasma samples from COVID-19 donors in the convalescent phase (3-6 weeks) were collected and subjected to comparative analyses among demographic factors (age, gender, and initial COVID-19 disease severity), antibody, and proteomic responses (**Fig. 1**) (**Table. S1**). Samples collected from healthy individuals collected pre-COVID-19 were included for comparison between health vs. convalescent COVID-19 (**Fig.1**). We first evaluated antibody responses detected in saliva and plasma specific for the SARS-CoV-2 receptor-binding domain (RBD), subunit 1 (S1), and subunit 2 (S2) of the spike protein, and the nucleoprotein (NP) (**Fig. 2A-C**). Antibodies to common cold coronaviruses were also evaluated by measuring antibodies that bind to the NL63 (NL63) Spike Glycoprotein (S1). Our primary interest was antibodies specific to the RBD and S1 of SARS-CoV-2, the sites responsible for virus binding to cell receptors and major mutation sites. When comparing antibody titers in convalescent subjects versus healthy controls, significant increases were observed in RBD binding IgA in saliva (p=0.0001), RBD binding IgA in plasma (p=0.0003), S1 RBD binding IgG in plasma (p<0.0001), and RBD binding IgM in saliva (p=0.0118) (**Fig. 2A-C**). Interestingly, significant correlations between paired saliva and plasma were also observed for SARS-CoV-2 RBD or S1 binding immunoglobulins (**Fig. 2D**). Significant correlations among immunoglobulin subclasses in plasma and saliva are summarized in **Table S2**.

**Fig. 1.**
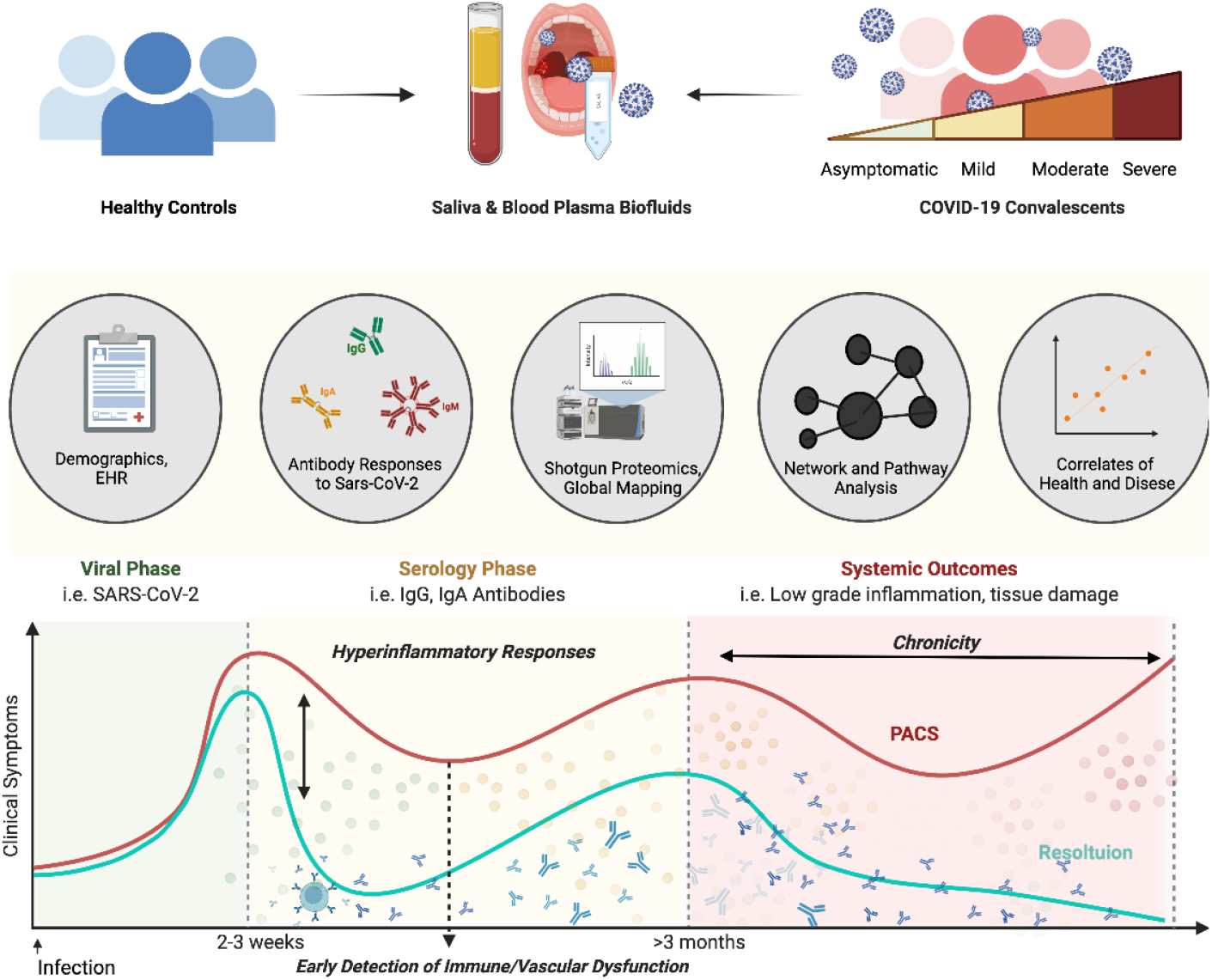
Study Design. Saliva and plasma samples were collected from convalescent coronavirus disease (COVID-19) donors (n=34) and healthy (n=13) to investigate the viral-immune axis in health versus disease. The serology coupled with the global shotgun proteomic analysis of plasma and saliva samples was conducted in parallel, followed by correlation analyses to demographic factors, antibody-, and proteomic responses. This study was designed to capture the inflammatory response (yellow/red dots)during the start of the convalescent phase (>2 weeks after clinical symptom; antibody drawings) and investigate the correlation between biological, and demographic factors. Ultimately, our findings will be applied to discover early detection markers for the post-acute sequelae of SARS-CoV-2 infection.

**Fig. 2.**
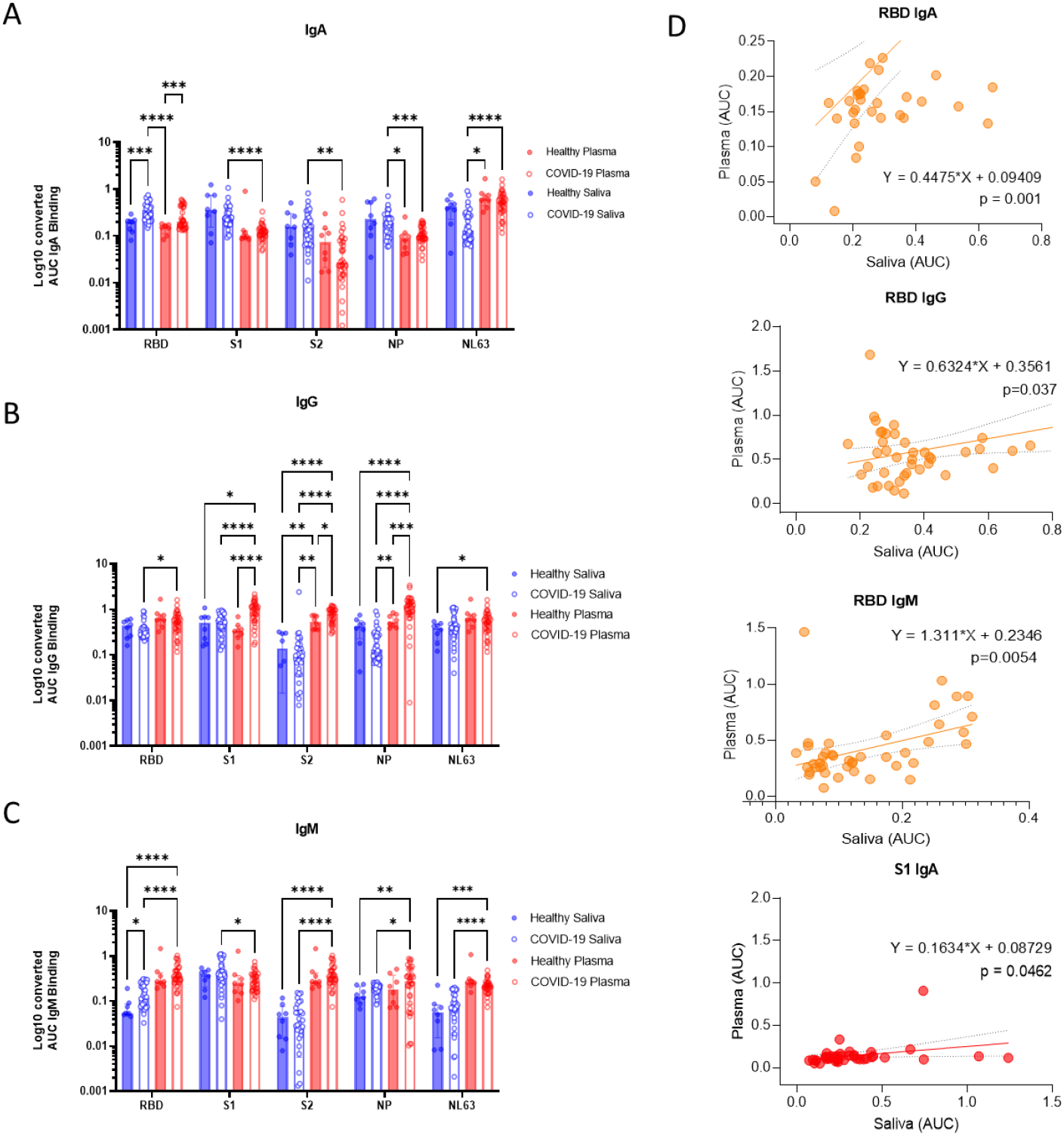
Compartmentalized antibody responses found in saliva and plasma collaborate in response to the SARS-CoV-2 infection. (A-C) The individual area under the curve (AUC) was plotted as blue or red hollow circles (saliva or plasma, respectively). Bars and whiskers represent median and standard deviation, respectively. Mixed-effect analysis with Tukey’s multiple comparisons test was used to measure statistical significance. *p ≤ 0.05; **p ≤ 0.01; ***p ≤ 0.001; ****p ≤ 0.0001. (D) Five paired immunoglobulins showing significant correlation (p<0.05) between plasma and saliva were depicted as simple linear regression models. The individual titer of saliva was plotted by paired plasma titers. The predicted regression line and deviations were depicted as a solid and dotted lines, respectively. Functions and p-value of regression analyses were indicated next to the regression lines. Correlations of immunoglobulins specific to the SARS-CoV-2 receptor binding site (RBD) or Spike protein 1 (S1) were colored yellow and red, respectively.

### Immunoglobulin composition and neutralizing functions displayed unique patterns between saliva and plasma

We next investigated the antibody responses to confirm that our subjects were in the COVID-19 convalescent phase and produced protective antibody responses. While antibody responses to the SARS-CoV-2 RBD and S1 showed a significant correlation between saliva and plasma, the overall antibody profile of saliva was different than plasma. For IgA response, convalescent saliva was significantly higher than convalescent plasma for SARS-CoV-2 S1, S2, NP, and NL63 (p<0.0001, p=0.0036 S1, p=0.0009 S2, and p<0.0001, respectively) (**Fig. 2A**). The IgG response showed an opposite trend in that the titers in convalescent plasma were significantly higher than in convalescent saliva for SARS-CoV-2 RBD, S1, S2, and NP (p=0.0178, p<0.0001, p<0.0001, and p<0.0001) (**Fig. 2B**). The IgM response was also significantly higher in plasma than saliva for all four SARS-CoV and NL63 antigens (p<0.0001 for RBD, p=0.0117 for S1, p<0.0001 for S2, p=0.0338 for NP, and p<0.0001 for NL63).

We evaluated saliva and plasma for neutralizing activity against SARS-CoV-2 S bearing pseudovirus particles (rVSV-GFPΔG*Spike). Saliva showed obviously lower neutralizing activity in comparison to plasma (**Fig. S1**). Neutralizing activity in plasma samples was surprisingly higher than expected. Despite the fact that the donors in the healthy group were collected pre-COVID-19 era and may have never encountered the SARS-CoV-2 virus, more than half of their plasma displayed cross reactivity with a significant level of neutralizing activity (62.5 %, median IC50=271.10). Convalescent COVID-19 subjects showed increased neutralizing activity in plasma for both positive rate and titer (92.16%, median IC_50_=317.30). In contrast, paired saliva samples were poor at neutralizing the pseudoviral particles, despite the robust RBD S1-binding IgA responses detected by ELISA (**Fig. 2A**). Only after purification and concentration of the IgAs in saliva (*16*), neutralizing activity was detected within a limited range from two convalescent COVID-19 salivary samples (16.22%, IC_50_=10.00).

### A global proteome analysis identified differentially expressed proteins

To characterize oral mucosal and systemic responses following SARS-CoV-2 infection more comprehensively we profiled saliva and plasma samples with mass spectrometry and proteomics. Dimension reduction by principal component analysis (PCA) showed a separation of convalescent COVID-19 donors from healthy controls for both saliva and plasma samples (**Fig. 3A&B**). Differentially expressed (DE) proteins between convalescent versus healthy samples are displayed as in volcano plots (**Fig. 3C&D**) and all significant observations are summarized in **Table 1**. The DE proteins, significantly enriched in convalescent saliva and plasma (fold change>2, p-value<0.05), are presented as in bar graphs (**Fig. 3E&F**). There were no DE proteins significantly enriched in healthy over convalescent COVID-19 (fold change <−2, p-value<0.05 in **Fig. 3C&E**). In saliva, convalescent COVID-19 samples showed a significant increase in expression of moesin (Uniprot accession number, AC=P26038, fold changes= 2.622, p-value<0.001), transmembrane protease serin D (AC=O60235,fold change=2.109, p-value=0.004), alpha-actin-1 (AC=P12814, fold changes= 2.450, p-value=0.004), nuclear transport factor 2 (AC=P61970, fold changes=2.251, p-value=0.0498), and serpin B13 (AC=Q9UIV8, fold changes= 2.293, p-value=0.001) (**Fig. 3C, E** and **Table. 1A**). The significantly enriched DEs in convalescent plasma in comparison to the healthy plasma were the fibrinogen alpha chain (AC=P0267, fold changes=3.279, p-value<0.001), Keratin, type II cytoskeletal 1 (AC=P04264, fold change=2.0231, p-value<0.001), apolipoprotein C-II (AC=P02655, fold changes=3.688, p=0.027), and REST corepressor2 (AC=Q8IZ40, fold changes=2.247, p-value=0.0122) (**Fig. 3D, F**, and **Table. 1B**). Significant proteins showed enrichment by acute COVID-19 disease severity (No symptom (n=15), mild (n=19), and moderate to severe (n=13)) including salivary IL-1 inhibitor, salivary sulfhydryl oxidase 1, salivary alpha-1B-glycoprotein, salivary protein kinase C inhibitor protein 1, salivary Immunoglobulin kappa chain V-III region CLL, plasma apolipoprotein A-IV, plasma REST corepressor 2, and plasma cytokeratin-1 (**Fig. 3F**). Salivary moesin and plasma fibrinogen alpha, which showed robust expression in participants who recovered from mild illness, clearly differentiated no-symptom participants but did not show a further increase in moderate to severe cases.

**Fig. 3.**
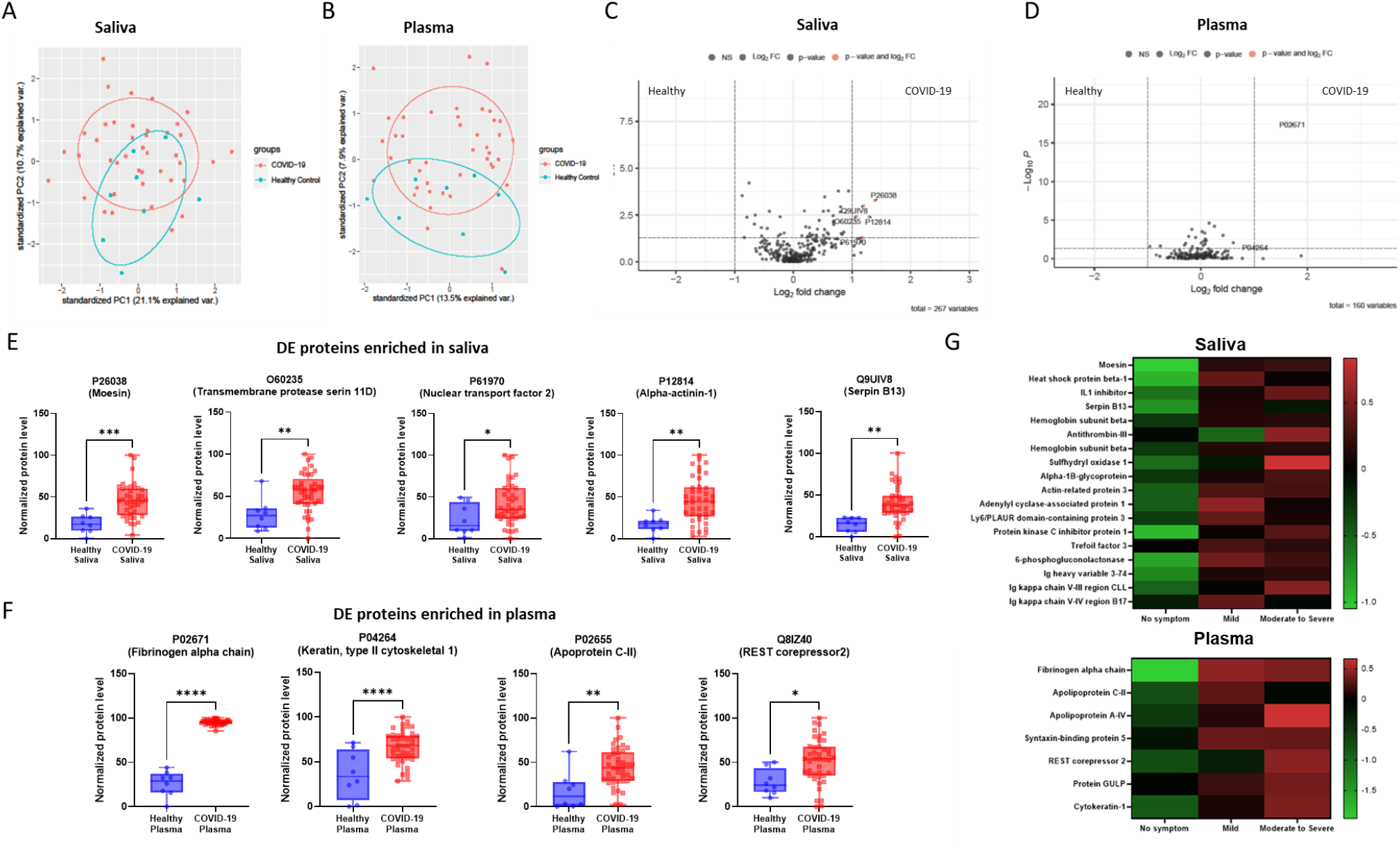
Comparative proteomic analyses revealed differentially expressed proteins (DEs) enriched in convalescent COVID-19 saliva and plasma. (A&B) Dimension reduction by principal component analysis (PCA) showed a separation of proteins from convalescent COVID-19 donors from healthy controls in saliva and plasma, respectively. Circles indicate 95% confidence intervals of group memberships. Percentages along the axes indicate the degree of variance explained by that principal component. (C&D) Fold changes of protein expression of convalescent COVID-19 over healthy samples were plotted by negative log-transformed p-values in volcano plots. Dotted lines of the volcano plot represent thresholds for the fold changes (Log2 fold changes >1) and statistical significance (p<0.05). (E&F) Differentially expressed proteins (Log2 fold changes>1, p<0.05) in saliva and plasma, were depicted as in a relative abundance of five significantly up-regulated proteins. Individual dots represent individual values. The interquartile range, median, and min/max values were illustrated as box, middle line, and whiskers, respectively. Mixed-effect analysis with Tukey’s multiple comparisons test was used to measure statistical significance. *p ≤ 0.05; **p ≤ 0.01; ***p ≤ 0.001; ****p ≤ 0.0001. (G) Heatmaps of proteins significantly enriched (p<0.05) in saliva (top) and plasma (bottom) were collected from moderate to severe participants, in comparison to no participants without apparent symptoms. Heatmap was color-coded based on the normalized level.

**Table 1.**
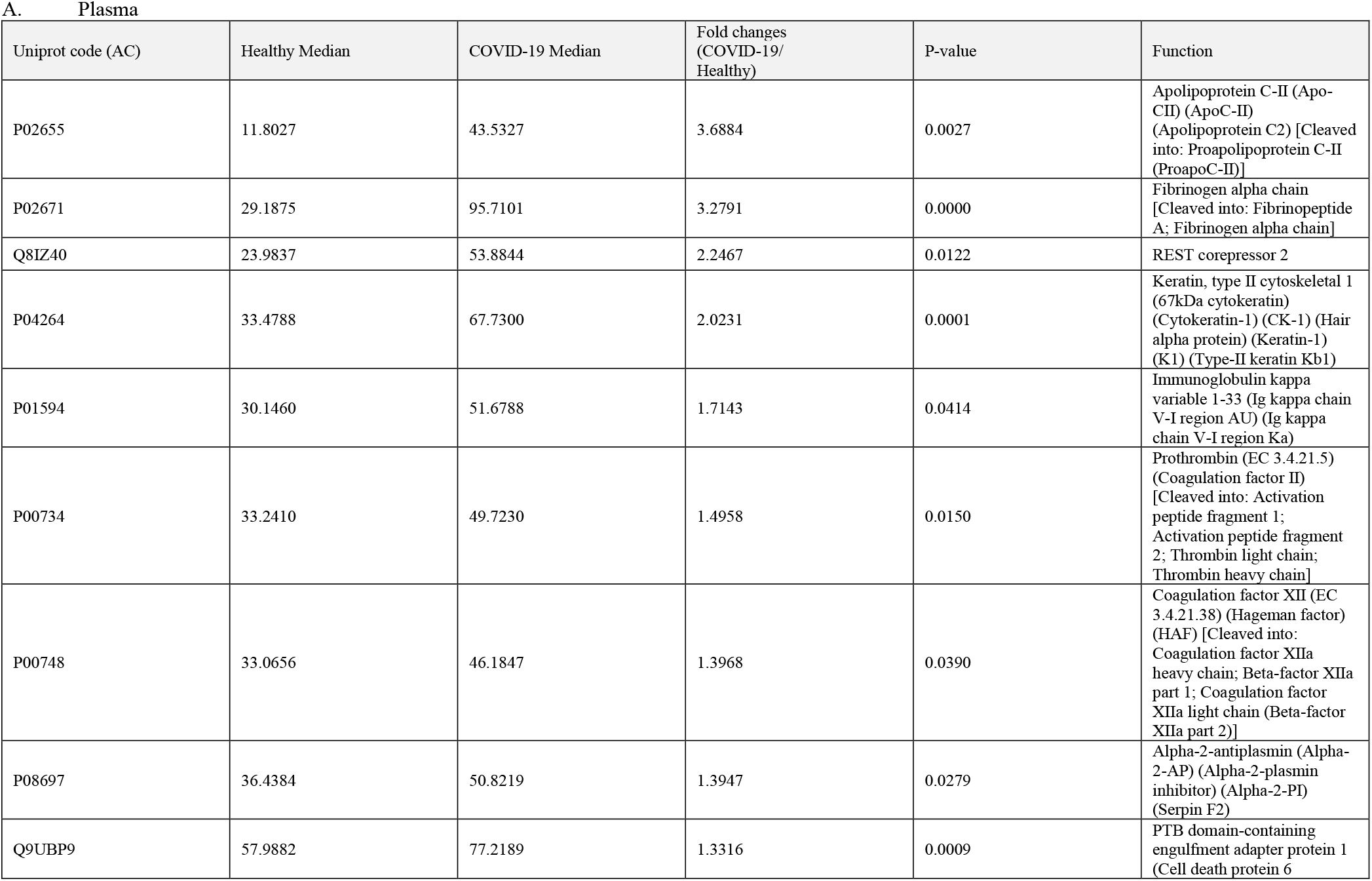
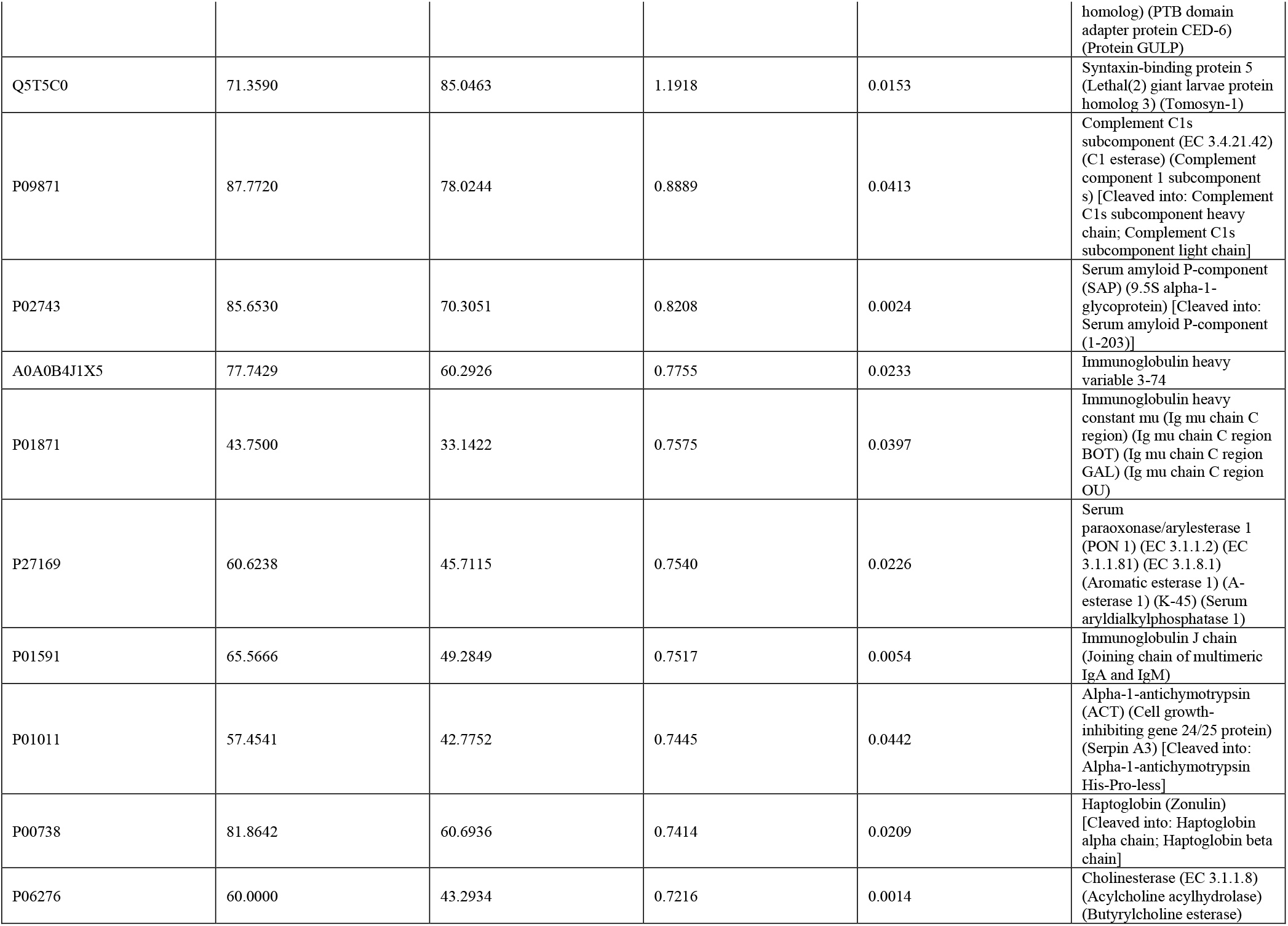
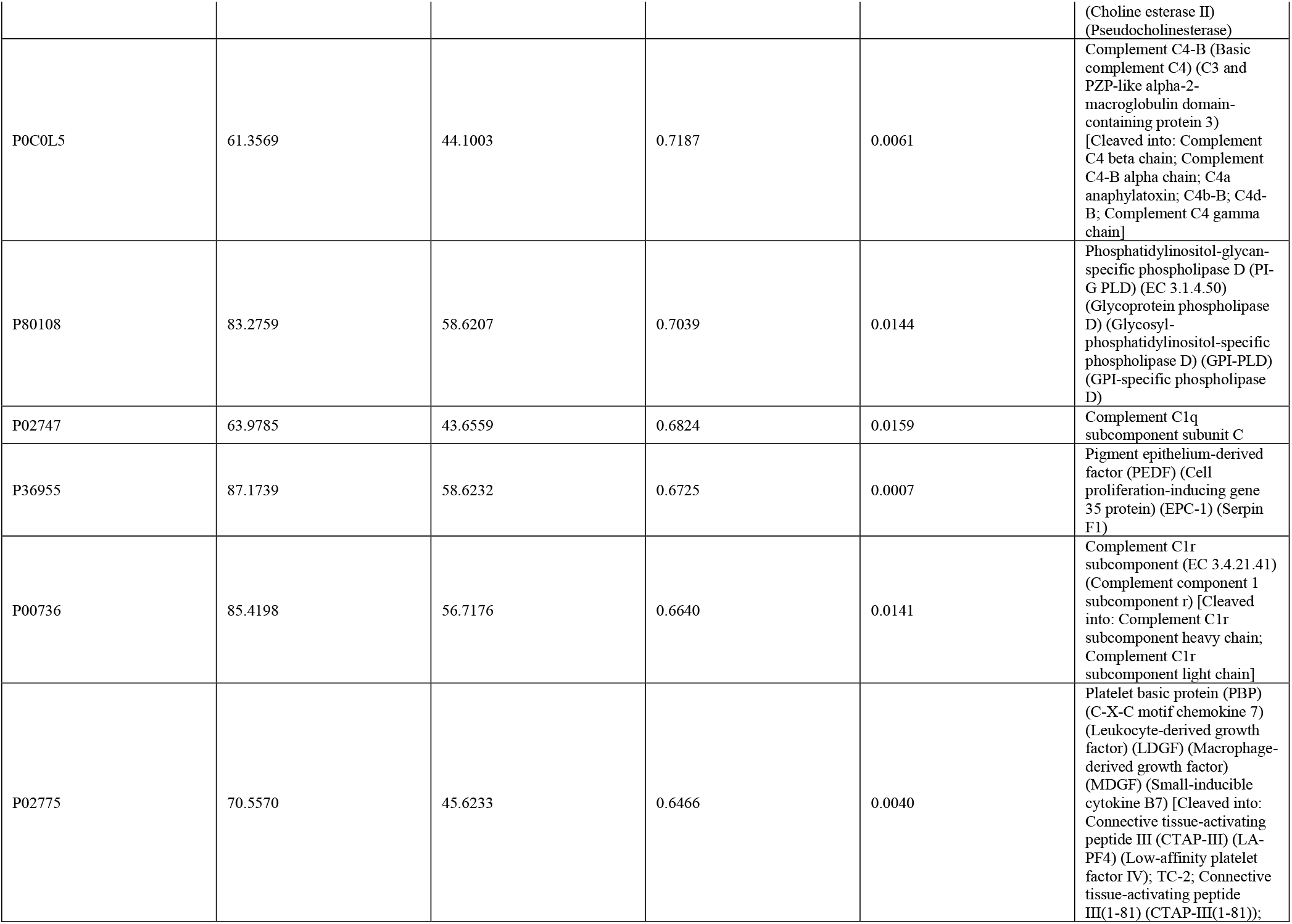
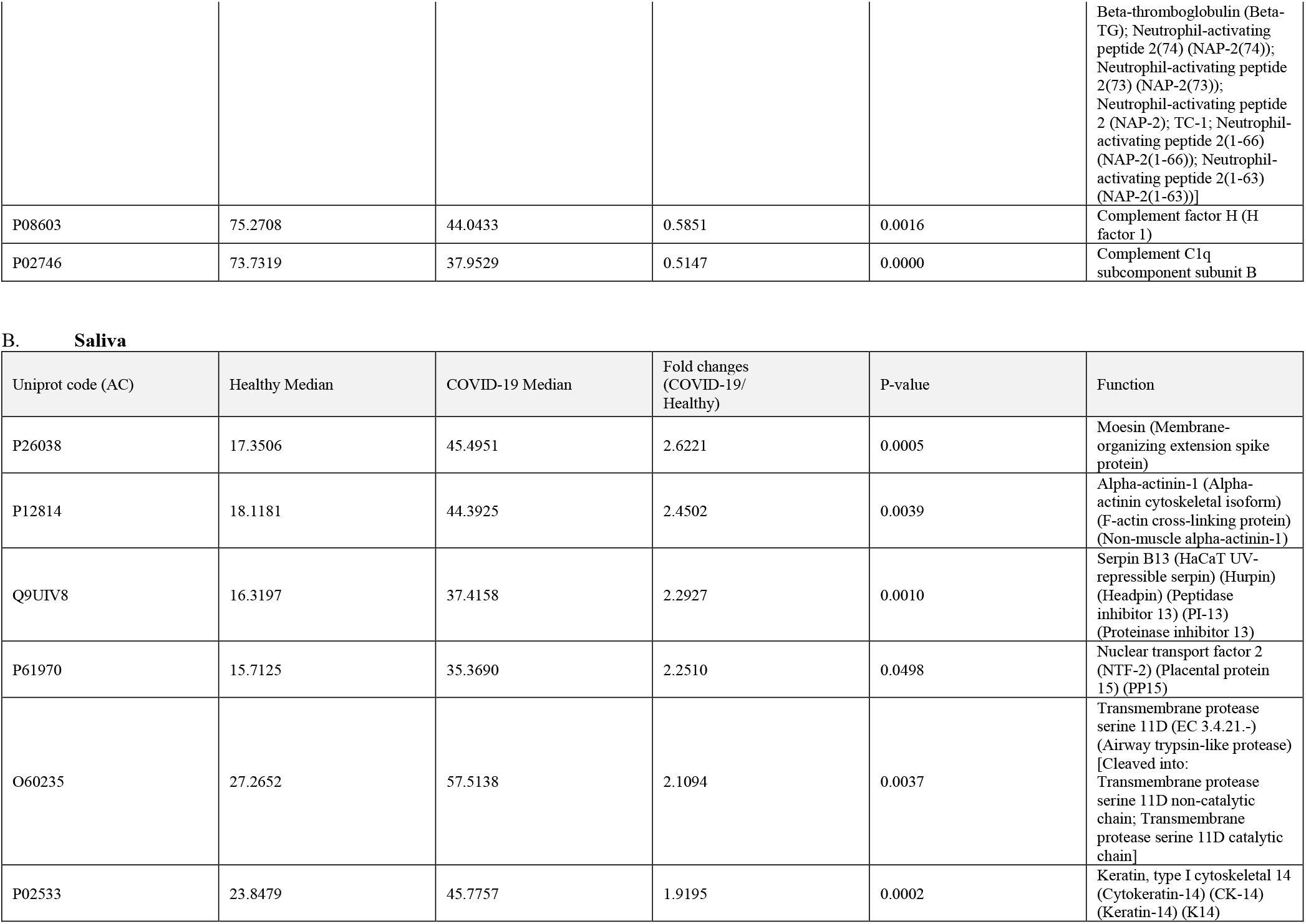
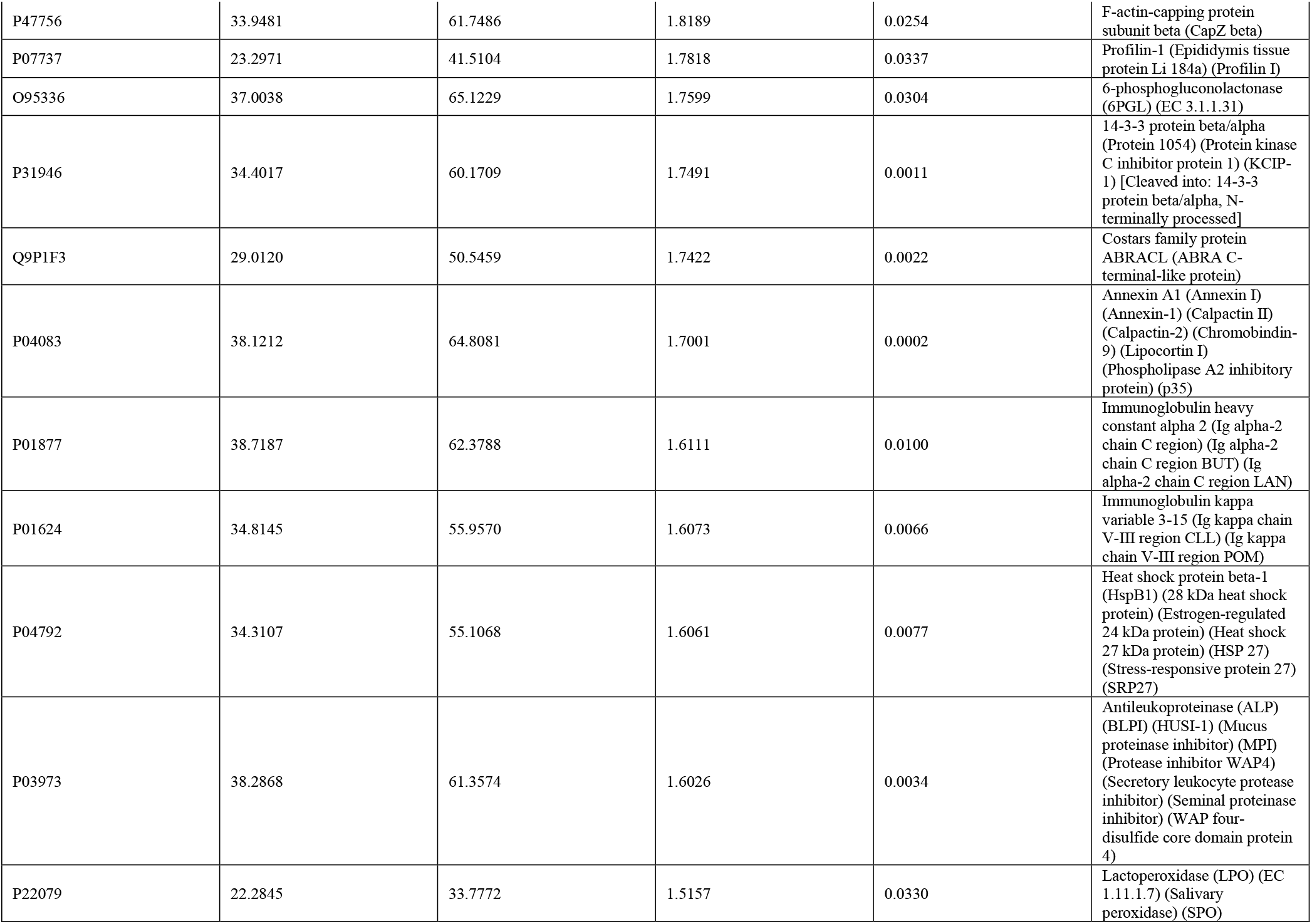
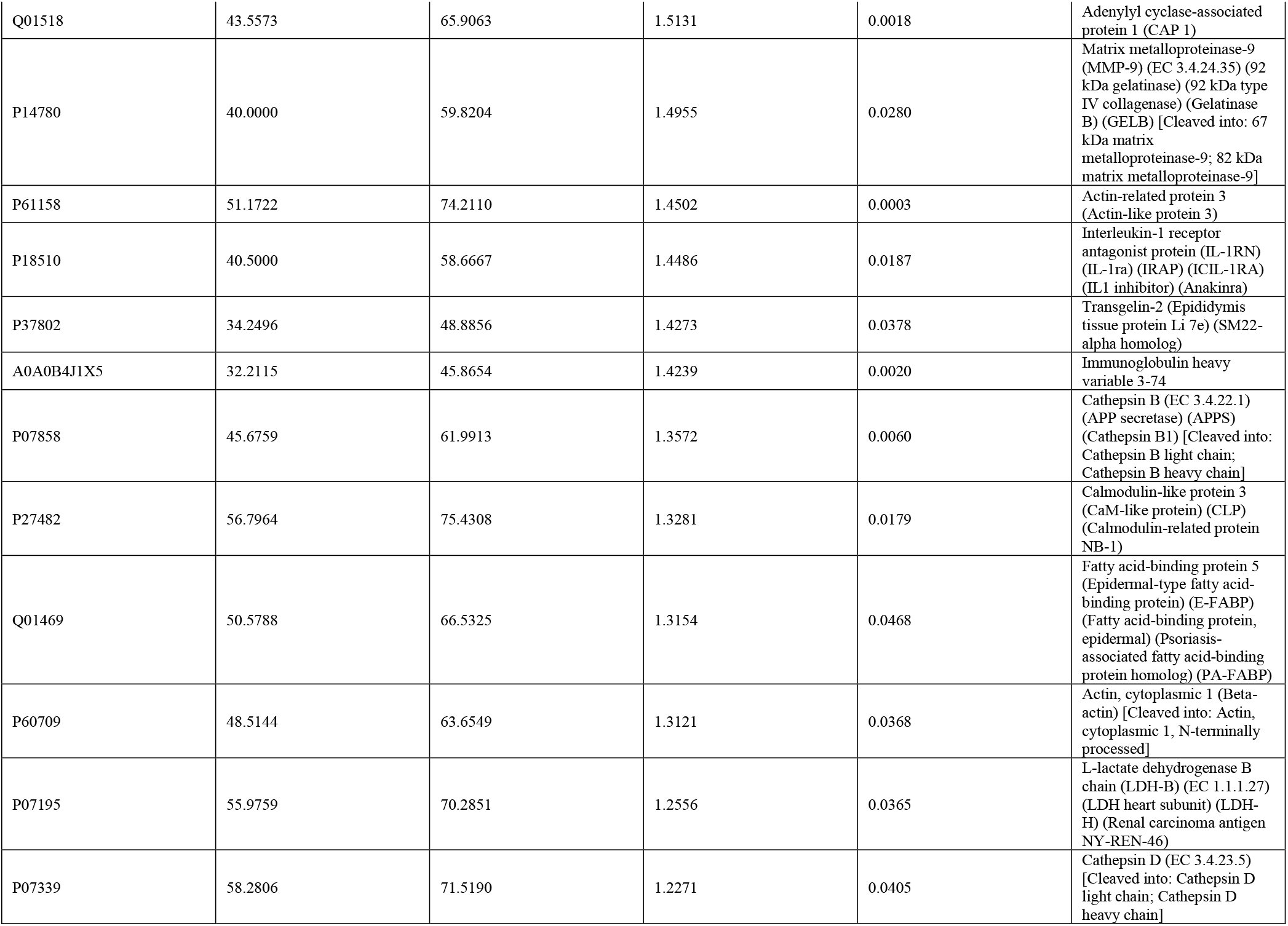
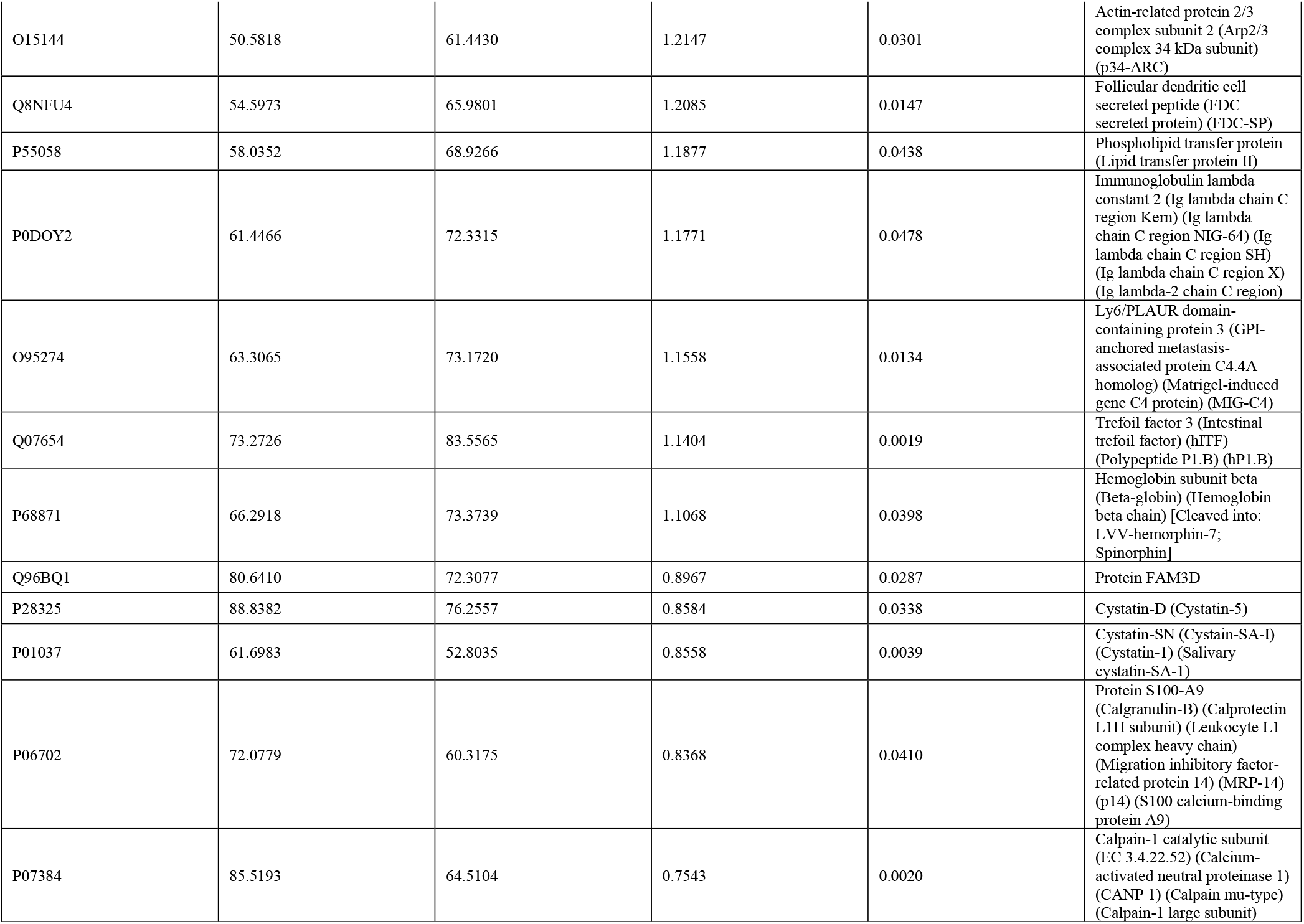
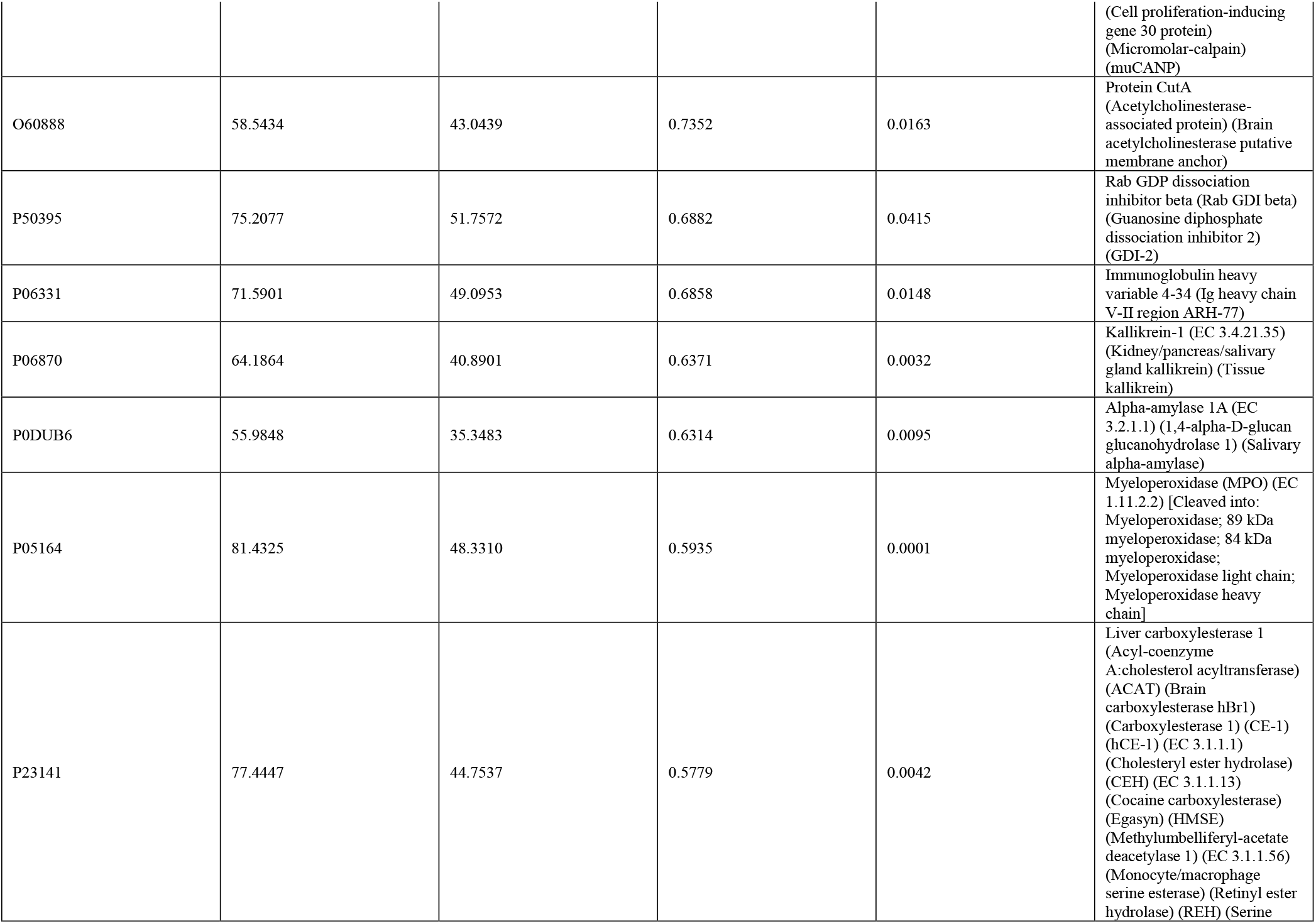
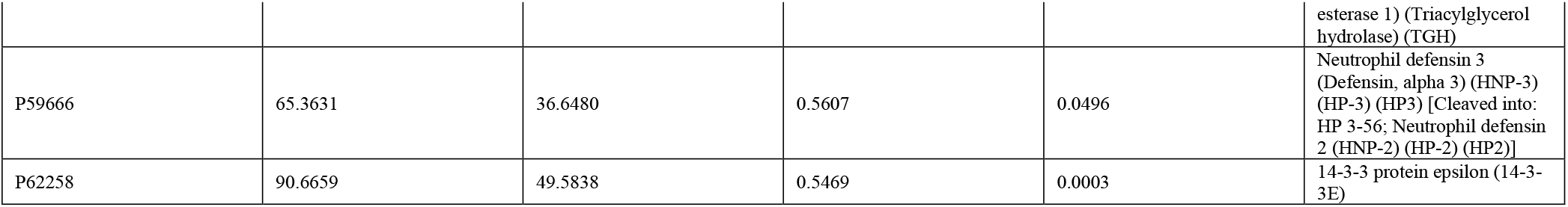
Significant observations (p<0.05) in fold changes of differentially expressed proteins in the saliva of convalescent COVID-19 and healthy samples.

The proteomic signature was evaluated by random forest machine learning and network analyses (STRING enrichment analyses). The hierarchical clustering heatmap generated from the random forest machine learning demonstrated the pathway is clustered into two groups based on convalescent COVID-19 vs. healthy (**Fig. 4A**). Network analyses performed based on differentially expressed proteins between convalescent COVID-19 and healthy samples (**Table 1**) showed that the convalescent plasma proteome displayed suppressed biological functions involved in oxidative damage response, and antimicrobial properties against opportunistic infection (*Staphylococcus aureus* infection) and complement and coagulation cascades (**Fig. 4B**). In contrast, the pathways enriched in convalescent COVID-19 plasma were all associated with fibrin clot formation (**Fig. 4B**). Pathways enriched in convalescent samples included hemostasis, platelet degranulation, immune system, interleukin-12 signaling, and leukocyte activation (**Fig. 4B**). Pathways related to granule or lysozyme formation were suppressed in saliva from convalescent COVID-19 donors (**Fig. 4B**).

**Fig. 4.**
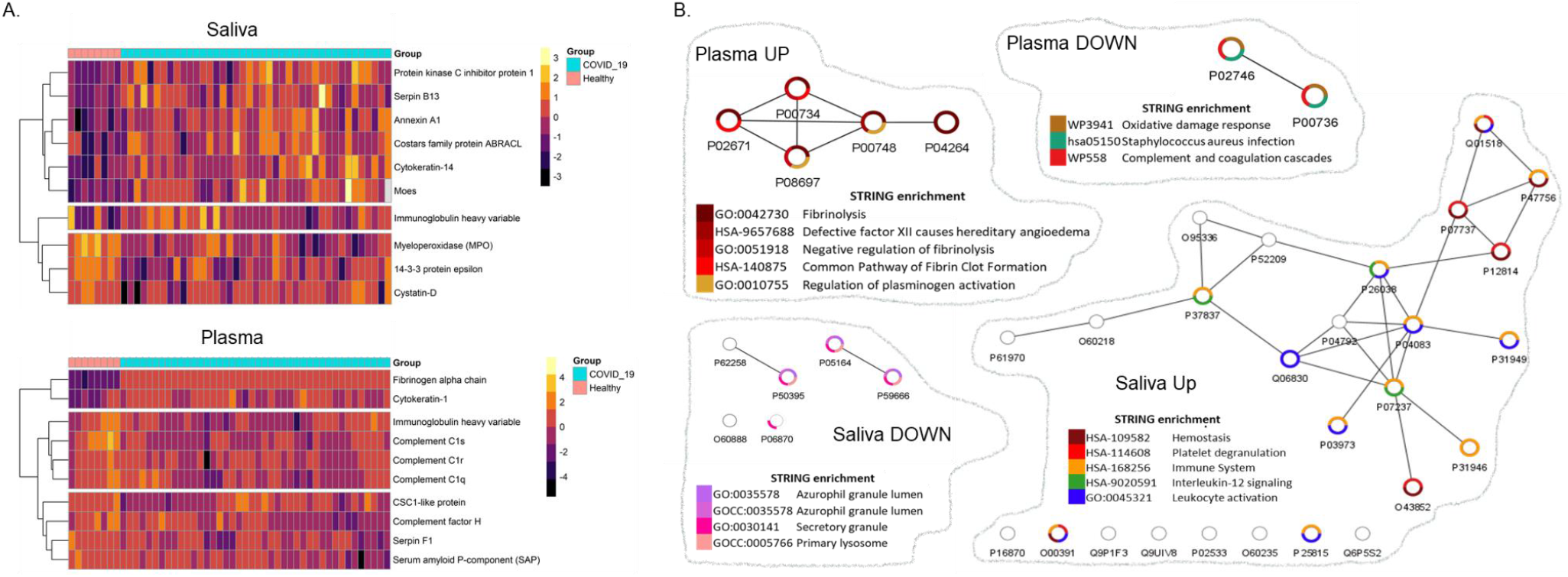
Network analyses depicted altered pathways and functions in convalescent plasma and saliva. (A) Heatmaps of proteins significantly enriched (p<0.05) in saliva (top) and plasma (bottom) collected from moderate to severe participants, in comparison to no participants without apparent symptoms. Heatmap was color-coded based on the normalized level. (B) Significantly enriched differentially expressed protein (DE) proteins in convalescent COVID-19 vs. Healthy individuals were subjected to the random forest machine learning and proteins that showed 100% accuracy predicted were plotted in the heatmap. A dendrogram was constructed based on hierarchical clustering and the group information (healthy vs. COVID-19) was color-coded. (C) In parallel, interactions among DE proteins were predicted by the STRING enrichment analyses and depicted as network maps using Cytoscape. Pathways associated with each DE protein were depicted in a donut graph, color-coded based on terms discovered by the STRING enrichment assay.

### Altered proteomic functions in COVID19 convalescent saliva directly correlated with the expression of RBD-binding IgA response in saliva

Comparative proteomic analyses between healthy vs. convalescent COVID-19 suggest that inflammatory markers induced by SARS-CoV-2 remain in both body fluids during the recovery phase. For a better understanding of the inflammatory patterns occurring in the oral local mucosal and systemic immune system, we performed a separate set of analyses that compared the salivary and plasma proteome (**Fig. 5A**). The PCA analysis showed a clear separation between the saliva and plasma proteome for both healthy and convalescent COVID-19 samples (**Fig.5B**). Hierarchical clustering heatmaps were clustered into two groups based on the origin of samples (saliva vs. plasma) while demographic factors (gender, age, and acute COVID-19 disease severity) did not contribute to the clustering. (**Fig. 5C**). In a healthy state, most DE proteins were higher in saliva, but proteins related to coagulation pathways (complement C3, C5, antithrombin) were higher in plasma (**Fig. 5C**). The convalescent COVID-19 heatmap had a similar pattern to the healthy heatmap, except that the expression pattern of apolipoprotein and fibrinogen was reversed (saliva>plasma in healthy; plasma>saliva in COVID-19, **Fig. 5C**). The cluster of saliva in the convalescent COVID-19 heatmap further diverged into three subclusters (Blue arrows and roman numerals on top of the **Fig. 5C**), suggesting a link between the humoral immune response and the innate inflammatory response in the oral mucosa. Each cluster was labeled as I, II, and III in increasing order of protein expression level.

**Fig. 5.**
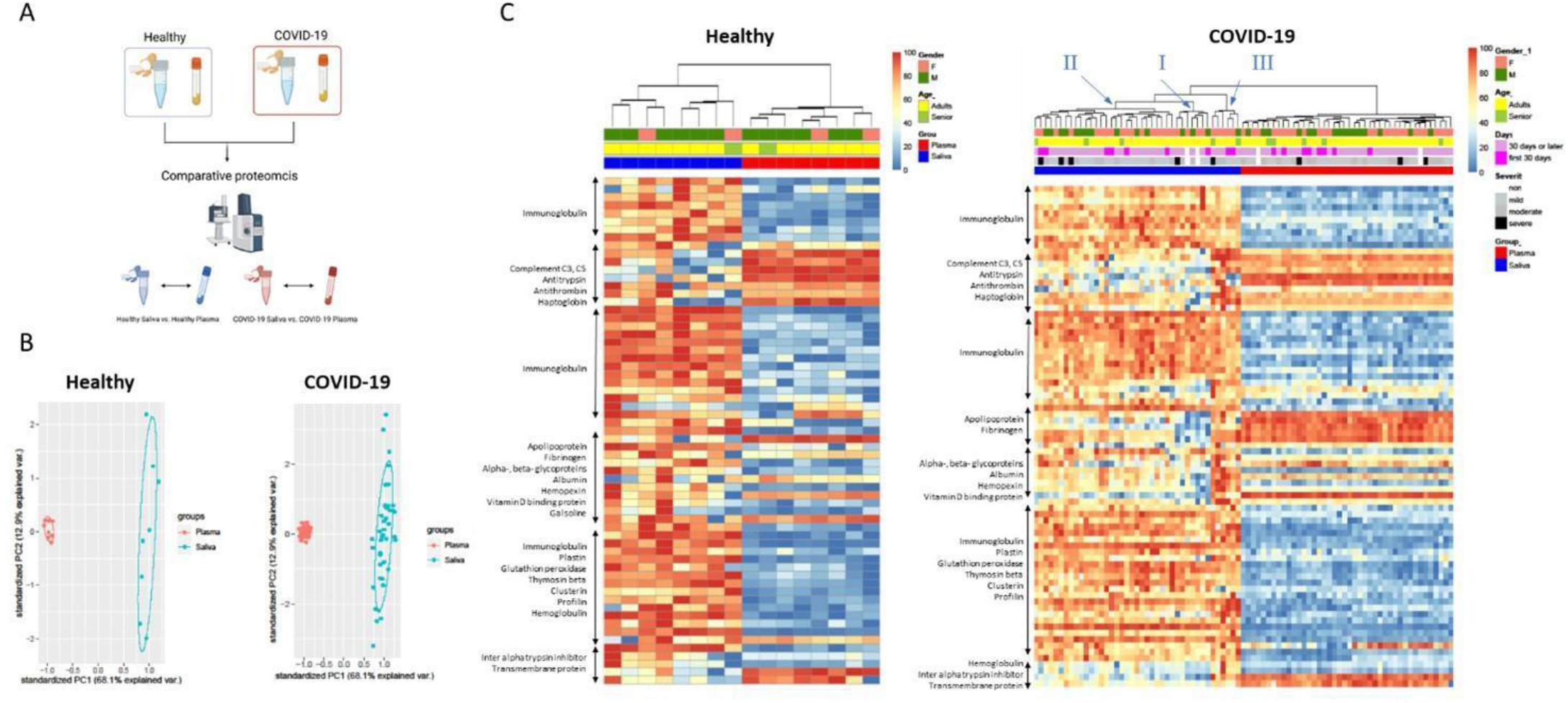
Comparative proteomic analyses between saliva and plasma revealed the heterologous signatures of each biofluid and further divarication of convalescent COVID-19 saliva. (A) The proteomics data were further analyzed to compare proteomic composition between saliva vs. plasma. The data obtained from healthy and convalescent COVID-19 participants were separately analyzed. (B) Differentially expressed proteins (Log2 fold changes>1, p<0.05) in saliva and plasma, were depicted as in a relative abundance of five significantly up-regulated proteins. The 95% confidence intervals of group memberships and degree of variance were indicated as circle and percentages on the axes, respectively. (C&D) The heterologous proteomic profile between saliva and plasma was further analyzed using clustered heatmap for both healthy and convalescent COVID-19. Relative abundance was calculated based on the proportion of normalized reads and displayed as color gradients. A dendrogram was constructed based on hierarchical clustering of relative abundances and color-coded demographic information of each participant was added to show their association with each clade. Blue Roman numerals and arrows indicate subclades in COVID-19 saliva. The numbering was in crescent with the expression level of DE proteins.

We then determined the influence of the immune subclusters in relation to serological results for each fluid by investigating correlations with immunoglobulin levels (**Fig.6A-C**). Strikingly, significant correlations were observed with the RBD-binding saliva IgA, IgM, and plasma IgA titers, as in a linear increase by the order of subcluster numbers (p=0.0274, p=0.0038, and p=0.0409, respectively, **Fig. 6A-C**). Since we observed an increasing trend in the expression level of DE protein, we also performed a correlation analysis between each RBD binding immunoglobulin with each DE protein involved in the convalescent COVID-19 saliva sub-clustering (**Fig. 6D-H**). The Clusterin showed significant positive correlations with RBD binding IgA in both saliva and plasma (**Fig. 6D&E**). Fibrinogen beta chain was significantly correlated with RBD binding IgA in saliva (**Fig. 6F**), and Apolipoprotein A1 was correlated with the RBD binding salivary IgM and plasma IgA (**Fig. 6G&H**).

**Fig. 6.**
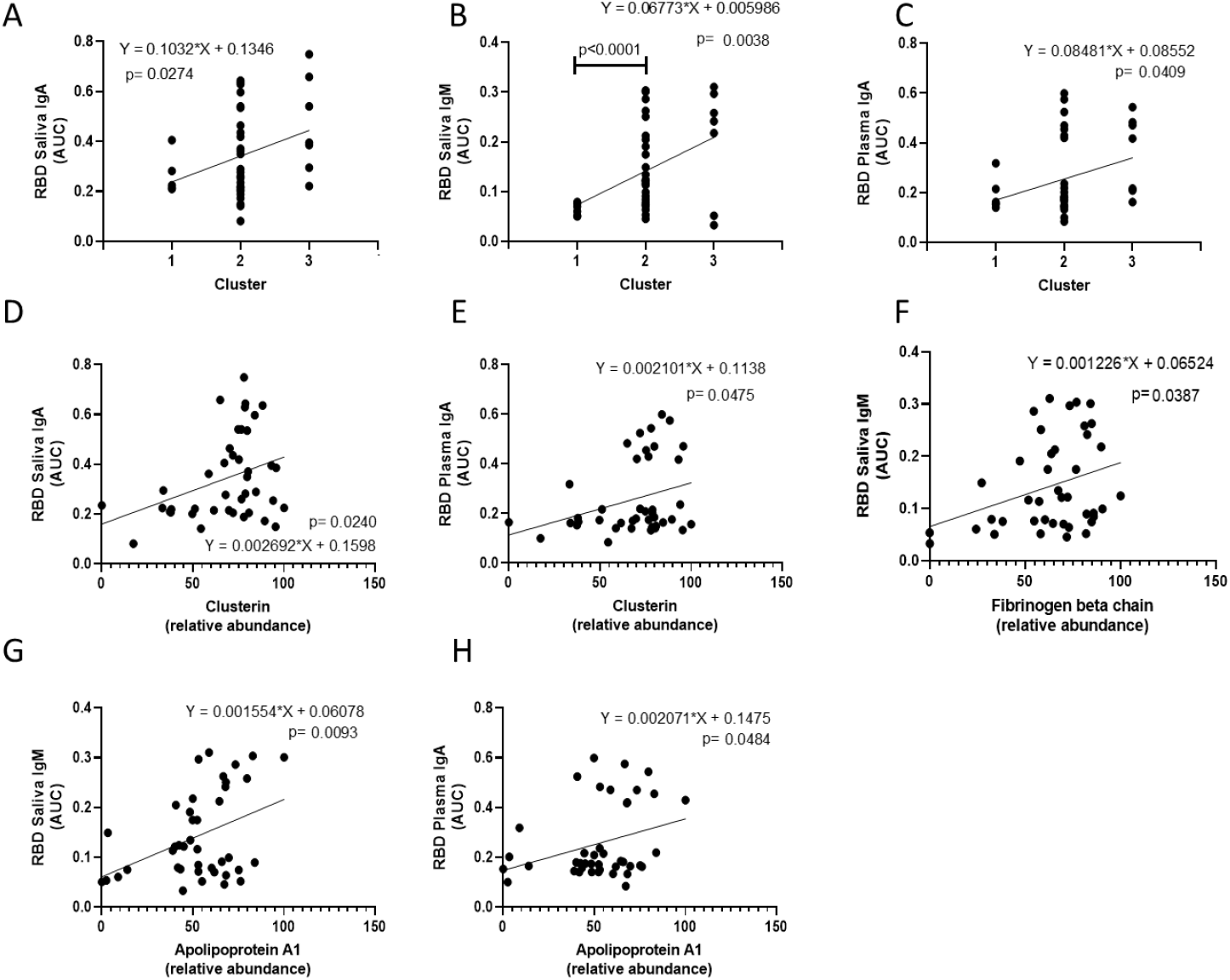
Correlation analyses suggests that proteomic alterations in convalescent saliva are associated with antibody responses specific to the receptor binding site (RBD) of SARS-CoV-2. (A-C) A significant correlation (p<0.05) between SARS-CoV-2 RBD specific immunoglobulins and convalescent COVID-19 salivary sub-clusters was depicted as simple linear regression models. The individual titer of immunoglobulin was plotted by subcluster numbers. The predicted regression line and deviations were depicted as solid and dotted lines, respectively. Functions and p-value of regression analyses were indicated next to the regression lines. (D-H) Three differentially expressed proteins responsible for the clustering were illustrated as simple regression models as described above.

Together, our results indicate that measurement of only antibody levels during the COVID-19 convalescent phase does not provide a full picture of the host response mounted after SARS-CoV-2 infection. Indeed, while we confirmed that our convalescent COVID-19 subjects had produced antibodies, our global proteomics analysis revealed novel aberrant immune signatures and clotting dynamics in plasma and saliva when compared to healthy controls. Overall, we demonstrated that population-based investigations of saliva can be used to map global host responses to local mucosal and systemic functions in addition to the characterization of antibody responses.

## Discussion

Increasing evidence indicates that the immune and endothelial health of convalescent COVID-19 subjects may be compromised when compared to healthy controls (*5*). Recovery from inflammatory responses to viral infection is mediated by multiple systems (*17*) and dependent on the overall host response. Molecular delineation of such components during physiological recovery versus pathologic transition is pivotal to develop host-directed strategies, aiming to sustain physiological function (*18*). Our findings also demonstrate that abnormal inflammatory and clotting responses can be identified in both saliva and plasma fluids of convalescent COVID-19 subjects. This indicates that even when SARS-CoV-2 antibody responses are mounted, the COVID-19 convalescent phase does not necessarily define disease resolution. We also highlight saliva as an important and accessible fluid that can be monitored to identify not just antibody responses, but also diverse immune pathways, including mucosal immunity, innate immune responses, neutrophil functions, and clotting pathways.

Carefully designed serosurveillance studies aimed at implementing antibody testing by investigations of blood-derived fluids (*19, 20*), but not saliva. Convalescent COVID-19 subjects from our study successfully mounted antibody responses to SARS-CoV-2 in both blood plasma and saliva fluids, confirming the clinical phase of our subjects and the feasibility of our investigations. While the majority of serological testing in SARS-CoV-2 cases detect IgG antibodies at individual and population levels (*21*), our study showed a significant increase in RBD binding IgA in both convalescent saliva and plasma, S1 binding IgG in plasma, and RBD binding IgM in saliva. Significant correlations between paired saliva and plasma showed positivity for SARS-CoV-2 RBD or S1 binding immunoglobulins, indicating that saliva is an available biofluid for monitoring the presence of protective antibody responses and immune responses. There were several similarities detected among both fluid types, we also found unique patterns within saliva when compared to blood plasma. The IgA response in convalescence was significantly higher in saliva than in plasma, whereas the IgG response showed an opposite trend in that the titers in convalescent plasma were significantly higher than in saliva. This is expected as IgG is the dominant subtype in the blood (*22*), while IgA is found in mucosal tissues (*23*). To date, however, evidence on antibody responses and neutralization levels to SARS-CoV-2 provided a limited range of information regarding the immune responses and pathogenesis of subjects that recovered, or not, from the natural infection.

Next-generation plasma profiling demonstrates a comprehensive overview of the immune response and has the potential to elucidate the impact of COVID-19 on the host. Zhong *et al.* showed that more than 200 proteins were found significantly different in plasma levels at the time of infection as compared to 14 days later (*24*). In comparison to Zhong et al’s findings, our plasma proteome appears to reflect a recovery process, displaying much fewer numbers of significantly enriched DE proteins (p-value<0.05, fold change>2) (Fig. 3D and Table 1). Yet, the participants of our cohort still displayed a significant enrichment of fibrinogen in plasma. If not limited to the proteins upregulated by 2 fold or higher, convalescent plasma showed an increase in numerous proteins associated with neutrophil functions or migration, such as annexin 1 (*25, 26*), antileukoproteinase (*27*), and Matrix metalloproteinase-9 (*28, 29*) (**Table 1A**). Interestingly, salivary proteome appears to maintain activated inflammatory status longer than plasma, as significantly increased neutrophil activation markers, myeloperoxidase (MPO), annexin 1-2, alpha-actinin-1, and nuclear transport factor 2 are involved in the migration of neutrophils (*30–32*). The convalescent saliva fluid also showed a significant increase in transmembrane protease serine 11D, which is known to activate the SARS-CoV-2 spike protein and facilitate the viral-cell fusion process (*33, 34*), and serpin B13, known for regulating neutrophil serine proteases and inflammatory caspases (*35*) (**Table 1B**). Other interesting proteins found in our study are the inhibitors of cathepsin, which have been involved in viral cell entry and replication (*36*). This was found to be significantly higher in plasma during acute infection versus convalescent COVID-19 cases and their analysis demonstrated that a group of patients display a “disease profile”, despite not having no symptoms of the disease (*24*).

We further demonstrated that salivary IgA antibody responses to SARS-COV-2 could be involved in neutrophil-fibrinogen interactions at the oral mucosal surface. The plasma fibrinogen alpha chain displayed a significant correlation with salivary markers, such as salivary RBD IgA and salivary IgM (**Table 2**). Significant correlations were also observed between the salivary RBD IgM and salivary RBD IgA; salivary RBD IgM and salivary fibrinogen beta chain; salivary RBD IgM and severity of clinical illness during the acute disease phase. This suggests that salivary antibodies to the SARS-CoV-2 infection participate in the inflammatory response mediated via neutrophil extracellular traps (NETs)-fibrin in oral mucosa and possibly contribute to the systemic inflammatory response represented by enhanced plasma fibrinogen. In severe COVID-19, neutrophil degradation and NETosis in blood and in the lung have been functionally linked to severe inflammation and thrombosis (*37*–*42*). Abnormal fibrinolysis is known to impact networks with neutrophil functions, including NET formation (*38, 43, 44*). Indeed, excessive release of NETs, with a low resolution of inflammation, can lead to immune thrombosis in blood vessels, with NET-fibrin interactions contributing to the severity of tissue injury and pathogenesis (*45*). Unique to the oral organ, the NET-fibrin axis also plays a unique role in regulating the constant deposition of fibrin produced by the commensal microbiome-triggered inflammation (*46*).

**Table 2.**
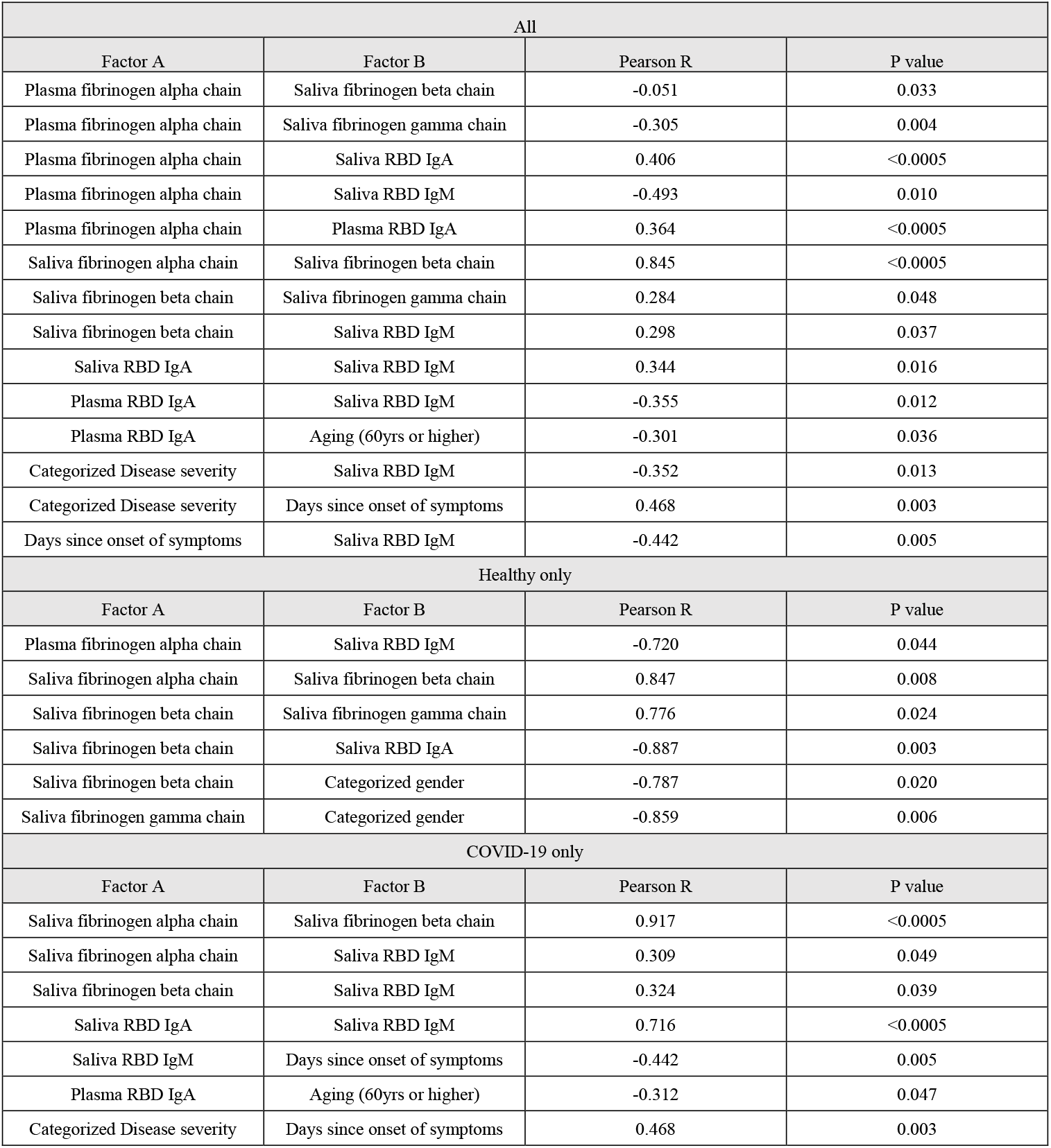
Correlations among RBD binding immunoglobulins, fibrinogen proteins, and demographic factors.

| All |  |  |  |
| --- | --- | --- | --- |
| Factor A | Factor B | Pearson R | P value |
| Plasma fibrinogen alpha chain | Saliva fibrinogen beta chain | -0.051 | 0.033 |
| Plasma fibrinogen alpha chain | Saliva fibrinogen gamma chain | -0.305 | 0.004 |
| Plasma fibrinogen alpha chain | Saliva RBD IgA | 0.406 | <0.0005 |
| Plasma fibrinogen alpha chain | Saliva RBD IgM | -0.493 | 0.010 |
| Plasma fibrinogen alpha chain | Plasma RBD IgA | 0.364 | <0.0005 |
| Saliva fibrinogen alpha chain | Saliva fibrinogen beta chain | 0.845 | <0.0005 |
| Saliva fibrinogen beta chain | Saliva fibrinogen gamma chain | 0.284 | 0.048 |
| Saliva fibrinogen beta chain | Saliva RBD IgM | 0.298 | 0.037 |
| Saliva RBD IgA | Saliva RBD IgM | 0.344 | 0.016 |
| Plasma RBD IgA | Saliva RBD IgM | -0.355 | 0.012 |
| Plasma RBD IgA | Aging (60yrs or higher) | -0.301 | 0.036 |
| Categorized Disease severity | Saliva RBD IgM | -0.352 | 0.013 |
| Categorized Disease severity | Days since onset of symptoms | 0.468 | 0.003 |
| Days since onset of symptoms | Saliva RBD IgM | -0.442 | 0.005 |
| Healthy only |  |  |  |
| Factor A | Factor B | Pearson R | P value |
| Plasma fibrinogen alpha chain | Saliva RBD IgM | -0.720 | 0.044 |
| Saliva fibrinogen alpha chain | Saliva fibrinogen beta chain | 0.847 | 0.008 |
| Saliva fibrinogen beta chain | Saliva fibrinogen gamma chain | 0.776 | 0.024 |
| Saliva fibrinogen beta chain | Saliva RBD IgA | -0.887 | 0.003 |
| Saliva fibrinogen beta chain | Categorized gender | -0.787 | 0.020 |
| Saliva fibrinogen gamma chain | Categorized gender | -0.859 | 0.006 |
| COVID-19 only |  |  |  |
| Factor A | Factor B | Pearson R | P value |
| Saliva fibrinogen alpha chain | Saliva fibrinogen beta chain | 0.917 | <0.0005 |
| Saliva fibrinogen alpha chain | Saliva RBD IgM | 0.309 | 0.049 |
| Saliva fibrinogen beta chain | Saliva RBD IgM | 0.324 | 0.039 |
| Saliva RBD IgA | Saliva RBD IgM | 0.716 | <0.0005 |
| Saliva RBD IgM | Days since onset of symptoms | -0.442 | 0.005 |
| Plasma RBD IgA | Aging (60yrs or higher) | -0.312 | 0.047 |
| Categorized Disease severity | Days since onset of symptoms | 0.468 | 0.003 |

Other specific drivers of the abnormal inflammatory and clotting responses observed in our convalescent COVID-19 subjects require further study. Several research teams have identified SARS-CoV-2 or protein in “viral reservoir” tissue samples collected from subjects months after acute COVID-19 (*47*, *48*). For example, Gaebler *et al.* identified SARS-CoV-2 RNA and protein in 7 of 14 intestinal tissue samples obtained from asymptomatic COVID-19 patients with negative nasal-swab PCR at an average of 4 months after acute disease (*49*). The SARS-CoV-2 spike antigen S1 itself appears capable of directly interacting with platelets and fibrinogen to drive blood hypercoagulation (*50*). This suggests that further studies of convalescent COVID-19 saliva and plasma would benefit from the measurement of SARS-CoV-2 RNA and spike antigen in addition to inflammatory and proteomic signatures. SARS-CoV-2 persistence in intestinal tissue or the oral mucosa, and possible shedding of spike antigen into saliva or blood, could also perpetuate chronic inflammatory and clotting sequelae.

The molecular mechanisms underlying higher concentrations of IgA but lower IgA neutralizing activity in convalescent saliva also require further exploration. It is possible that higher salivary IgA concentrations represent some form of extended antibody-mediated disease enhancement. Antibody-mediated disease enhancement has been reported in diverse RNA viral diseases, such as influenza, SARS-CoV-2, Dengue, and human immunodeficiency viruses infections (*51–54*). One team found that SARS-CoV-2 RBD-specific neutralizing dimeric IgAs isolated from nasal turbinate could facilitate viral infection, transmission, and injury in Syrian hamsters (*55*). Aleyd *et al* (*56*) demonstrated that IgA enhances NETosis as an effective defense mechanism to eliminate pathogens at mucosal surfaces. In contrast, neutrophil activation by IgA immune complex is also known to contribute to the immunopathogenesis of autoimmune diseases, such as IgA vasculitis, and nephropathy (*57–59*). In respiratory viral disease models, such as influenza and SARS-CoV-2, the formation of an IgA-virus immune complex led to exacerbated NETosis of neutrophils isolated from peripheral blood mononuclear cells (PBMCs) *ex vivo* (*60*).

Our study has several limitations. Samples were collected at only a one-time point, and antibody levels or proteomic responses were not adjusted by the different baseline of each individual intervariability. It was also not possible to draw predictive conclusions from our findings but instead predictive correlations. While study subjects were able to report the severity of their acute COVID-19 illness (asymptomatic, mild, or moderate/severe), clinical symptom data was not obtained after convalescent phase when saliva and plasma were collected.

Future studies would benefit from requiring convalescent COVID-19 subjects to report possible chronic symptoms longitudinally. This is especially pressing since up to 30% of patients infected with SARS-CoV-2 are developing a wide range of persistent symptoms that do not resolve over months or years (*61*). These patients are being given the diagnosis LongCovid or Post-Acute Sequelae of COVID-19 (PASC) (*62*). Persistence of SARS-CoV-2 in tissue, aberrant immune signaling, and microclot formation have been documented in PASC (*63, 64*), but early molecular signs indicating direct risk to chronic symptoms have been elusive. Our current study sets the stage for global immune and proteome analyses that characterize inflammatory and clotting processes in PACs saliva and plasma in a manner that may be able to elucidate key aspects of the disease process and contribute to the development of targeted therapeutics.

## Materials and Methods

The research reported in this manuscript complies with all relevant ethical regulations and informed consent was obtained from all human participants. Additional information was collected on donor demographics (age and gender).

### Ethics Approval

This study has been approved by the University of California, San Diego Institutional Review Board, and the J. Craig Venter Institute (IRB, no. 200236X) and the J. Craig Venter Institute (IRB, no. 2020-286).

### Experimental study design

Blood and saliva samples were collected from convalescent COVID-19 donors who visited the COVID clinic at the University of California, San Diego (n=34). Confirmed COVID-19 cases were defined as previously described (*65*). Throughout the sample collection, the major SARS-CoV-2 strain circulating throughout the study was the original strain (USA-WA1/2020) and the vaccine against SARS-CoV-2 was not available. For comparison, we included healthy donors (n=13) from the pre-pandemic era, and subjects recruited to the study signed the institutional review board (IRB)– approved consent form (# 2018-268) (*66*).

Peripheral blood samples were collected by venipuncture and collected into BD vacutainer SST tubes (Vitality Medical, Salt Lake City, Utah). After 1hr., the collected blood sample was centrifuged for serum separation. Saliva was collected by the “passive drool technique” using the Saliva Collection Aid (Salimetrics, Carlsbad, CA). All samples were aliquoted and stored at −80°C for long-term storage.

The general experimental approach was summarized in Figure 1. Briefly, all collected plasma and saliva samples were tested for SARS-CoV-2 specific antibodies by enzyme-linked immunosorbent assay (ELISA) and pseudovirus neutralization assay (**Fig. 2 and Suppl. Fig. 1**, respectively). Correlations among all immunoglobulins (Ig) were investigated with Pearson’s correlation and simple linear regression analysis using the GraphPad Prism version 8.3.1. In parallel, separate sets of samples were processed and used for mass spectrometry to detect host antiviral-, and microbial proteins and peptides (**Fig. 3-5**). In the end, all collected data were collectively analyzed to verify the interaction among systemic and oral mucosal immune responses to the SARS-CoV-2 infection (**Fig. 6**).

### Antibody responses

#### SARS-CoV-2 binding ELISAs

Plasma and saliva samples were tested for binding to recombinant SARS-CoV-2 using an Enzyme-linked immunosorbent assay (ELISA) according to the manufacturers with slight modifications. The recombinant spike protein from human coronavirus NL63 (NL63) was also included as a coating antigen to estimate the presence of cross-reactive antibodies to common cold coronaviruses. All procedures were repeated twice, once manually and once by using Hamilton Microlab STAR (Hamilton, Reno, NV). Briefly, ELISA plates (Nunc MaxiSorp™ flat-bottom, Thermo Fisher Scientific, Waltham, MA) were coated with antigen (10ng/50μL) at 4 °C overnight. Four different coating antigens were included; SARS-CoV-2 Spike Glycoprotein (S1) RBD, His-Tag (HEK293)(NativeAntigen, Oxfordshore, United Kingdom), SARS-CoV-2 Spike Glycoprotein (S1), His-Tag (Insect Cells) (NativeAntigen, Oxfordshore, United Kingdom), SARS-CoV-2 Spike Glycoprotein (S2), His-Tag (Insect Cells) (NativeAntigen, Oxfordshore, United Kingdom), SARS-CoV-2 Nucleoprotein, His-Tag (E. coli) (NativeAntigen, Oxfordshore, United Kingdom), and Human Coronavirus NL63 Spike Glycoprotein (S1) His-Tag (HEK293) (NativeAntigen, Oxfordshore, United Kingdom). The next day, coated plates were washed three times with PBS-Tween (0.05%) and blocked with 200 μl of 5% milk blocking solution at room temperature for 30 min. During incubation, plasma and saliva samples were initially diluted 1:54 and 1:2, respectively, and three-fold serial dilution was performed. After blocking, diluted samples were added to the wells (50 μL/well) and incubated for 1 hr at room temperature. After 4X washing, 100 μL of 1:5000 diluted Goat anti-human secondary antibody was added into each well and incubated at 37°C for 1 hr (Goat Anti-Human IgG γ Chain Specific HRP conjugated, species Adsorbed (Human IgM, IgD, and IgA) Polyclonal Antibody for IgG (Cat# AP504P, EMD Millipore, Burlington, MA), Goat Anti-Human IgA, a-chain specific Peroxidase conjugate for IgA (Cat# 401132-2ML, Calbiochem, San Diego, CA) for IgA, and Goat Anti-Human IgM Fc5μ Fragment specific HRP conjugated secondary antibody for IgM (Cat# AP114P, EMD Millipore, Burlington, MA). After the incubation, plates were washed four times and 200 μL of the substrate (cat# P9187, Sigma, St. Louis, MO) was added for color development. After incubation in a dark room for 20 minutes, the reaction was stopped by the addition of 50 μL 3M H2SO4, and plates were read at 450 nm. The positive response was determined by the area under the curve (AUC) using GraphPad Prism version 8.3.1 (GraphPad Software, Inc., San Diego, CA, USA).

### Generation of pseudo-virus (rVSV-GFPΔG*Spike)

For pseudoviruses construction, spike genes from strain Wuhan-Hu-1 (GenBank: MN908947) were codon-optimized for human cells and cloned into eukaryotic expression plasmid pCAGGS to generate the envelope recombinant plasmids pCAGGS.S as described previously with slight modifications (*67*). For this VSV pseudovirus system, the backbone was provided by VSV G pseudotyped virus (G*ΔG-VSV) that packages expression cassettes for firefly luciferase instead of VSV-G in the VSV genome. Briefly, 293T cells were transfected with pCAGGS.S (30 μg for a T75 flask) using Lipofectamine 3000 (Invitrogen, L3000015) following the manufacturer’s instructions. Twenty-four hours later, the transfected cells were infected with G*ΔG-VSV with a multiplicity of four. Two hours after infection, cells were washed with PBS three times, and then a new complete culture medium was added. Twenty-four hours post-infection, SARS-CoV-2 pseudoviruses containing culture supernatants were harvested, filtered (0.45-μm pore size, Millipore, SLHP033RB), and stored at −70°C in 2-ml aliquots until use. The 50% tissue culture infectious dose (TCID50) of SARS-CoV-2 pseudovirus was determined using a single-use aliquot from the pseudovirus bank; all stocks were used only once to avoid inconsistencies that could have resulted from repeated freezing-thawing cycles. For titration of the SARS-CoV-2 pseudovirus, a 2-fold initial dilution was made in hexaplicate wells of 96-well culture plates followed by serial 3-fold dilutions (nine dilutions in total). The last column served as the cell control without the addition of pseudovirus. Then, the 96-well plates were seeded with trypsin-treated mammalian cells adjusted to a pre-defined concentration. After 24 h incubation in a 5% CO2 environment at 37°C, the culture supernatant was aspirated gently to leave 100 μl in each well; then, 100 μl of luciferase substrate (Perkinelmer, 6066769) was added to each well. Two min after incubation at room temperature, 150 μl of lysate was transferred to white solid 96-well plates for the detection of luminescence using a microplate luminometer (PerkinElmer, Ensight). The positive well was determined as ten-fold relative luminescence unit (RLU) values higher than the cell background. The 50% tissue culture infectious dose (TCID50) was calculated using the Reed–Muench method, as described previously (*68*).

### Pseudovirus neutralization assay

Neutralizing activity against rVSV-GFPΔG*Spike was determined as previously described with slight modification (*69*). Briefly, Vero cells were seeded at a density of 2.5 × 10^4^/50 μL in a Greiner Bio-One™ CellStar™μClear™ 96-Well, Cell Culture-Treated, Flat-Bottom, Half-Area Microplate (Thermo Fisher Scientific, Waltham, MA). The next day, the cell monolayer was rinsed with 0.01M PBS (Thermo Fisher Scientific, Waltham, MA). Due to the contamination by the commensal bacteria in saliva, total IgA was purified from saliva using Peptide M/agarose (InvivoGen Inc., San Diego, CA, USA) and used for the neutralization at low-dilution (1:2-1:10), as previously described(*16*). Plasma and saliva IgAs were three-fold diluted (starting from 1:50 and 1:2 dilution, respectively) with infection media (DMEM medium (cat# 11995065, Thermo Fisher Scientific, Waltham, MA) containing 2% fetal bovine serum (FBS)). Twenty-five μL of diluted samples was incubated with the same volume of pseudovirus (rVSV-GFPΔG*Spike) at 37°C for one hr. The sample-virus mixture was added to the Vero Cell monolayer and incubated at 37°C with 5% CO_2_. On the following day (12~16 hrs.), the expression of GFP was visualized and quantified by Celigo Image Cytometer (Cyntellect Inc, San Diego, CA). The neutralizing activity of the plasma sample was determined as pNT50 calculated from a transformed non-linear regression curve generated by GraphPad Prism version 8.3.1. (GraphPad Software, Inc., San Diego, CA, USA). Due to the low titer of salivary samples, the 50% inhibitory dilution (IC_50_) was determined by the reciprocal of the highest dilution of the sample corresponding to 50% reduction in GFP count compared with virus control minus sample control using the Reed-Muench method (*70*).

### Proteomics and peptidomics sample preparation

Deactivated saliva and plasma specimens were first passed through 10-kDa cutoff filters (Microcon, Millipore). The filtrates and the remaining materials on filters were subjected to peptidomics and proteomics analysis, respectively. For peptidomics analysis, the filtrates were dried in SpeedVac, and resuspended in 20 ul LC buffer A (0.1% formic acid in water). For proteomics analysis, the proteins remaining on filters were digested using the filter aided sample preparation (FASP) approach as described previously (*66*).

### Liquid Chromatography with tandem mass spectrometry (LC-MS/MS) analysis

For the LC-MS/MS analysis, the Ultimate 3000 nanoLC coupled to Q Exactive mass spectrometer (Thermo Scientific) was used as previously described (*9*). Peptides were first loaded onto a trap column (PepMap C18, 2 cm × 100 mm × I.D.; Thermo Scientific), and they were separated using an in-house packed analytical column (C18 ReproSil, 3.0 mm, Dr. Maisch GmbH; 20 cm x 75 mm I.D.) and binary buffer system (buffer A: 0.1% formic acid in water; buffer B: 0.1% formic acid in acetonitrile) with a 150-min gradient (2-35% buffer B over 105min; 35-80% buffer B over 10min; back to 2% B in 5 min for equilibration after staying on 80% B for 5 min). For the MS data acquisition, a top-10 data-dependent acquisition (DDA) method was applied. The maximum injection time was set to 20 ms, and the scan range was set to 350–1800 m/z with an AGC target of 1e6. The MS/MS acquisition was performed with 30% HCD collision energy. The target value was set to 5e5, and the maximum injection time was set to 100ms. Full MS and MS/MS scans were acquired at resolutions of 70,000 and 17,500, respectively. Dynamic exclusion was set to 20s. The mass to charge ratio (m/z [Da]) from mass spectrometry data was normalized and used for the calculation of fold changes of differentially expressed (DE) proteins (health vs. COVID-19; saliva vs. plasma).

### Database Search and Bioinformatics Analysis

For proteomics data analysis, protein identification and quantitation were performed using the MaxQuant-Andromeda software suite (version 1.6.3.4) as previously described(*71*)^8^. The majority of the default settings were taken, including trypsin as the enzyme, two missed cleavage sites, peptide length with minimum of seven amino acids, oxidation (M) as variable modification, and carbamidomethylation (C) as fixed modification. A UniProt human sequence database (20,376 sequences) was used for the protein database search. The false discovery rate (FDR) was set at 1% on both protein and peptide levels. Significantly enriched proteins in convalescent samples (p<0.05)(**Table 2**) were subjected to the network analysis by STRING enrichment analysis (Cytoscape software v. 3.9.1)(*72*). The heatmap was created using the pheatmap package in R using a hierarchical distance matrix and clustering option (*73*). The volcano plots were generated using the EnhancedVolcano package in R (*73*).

### Statistics

Data was statistically analyzed using the R or Graphpad Prism-8 suites of software (GraphPad Software, Inc., San Diego, CA, USA). Representatives of a minimum of two independent experiments were presented as the median and standard deviation and are representative of a minimum of two independent experiments. Data points for quantitative *in vitro* experiments represent all technical repeats for experiments done in triplicate. Antibody titers were analyzed by mixed-effect analysis with Tukey’s multiple comparisons.The significance of fold changes in DE proteins were measured using the Student’s t-test. Correlation among different parameters (antibody titers, proteomic marker expression levels, categorized demographic information, and salivary protein subgrouping) was evaluated by both Pearson’s R and simple linear regression analyses using Graphpad Prism-8 suites of software (GraphPad Software, Inc., San Diego, CA, USA).

## Acknowledgments

We would like to thank Wan Choi for his technical support, and staff at the J. Craig Venter Institute and University of California San Diego for support during the sample collection and transferring.

## Funding

The Conrad Prebys Foundation Grant (20-122) (MF, GT).

U.S. Public Health Service Grants (R00 DE0234804) from the National Institute of Dental and Cranial Research (MF).

## Author contributions

Conceptualization: MF

Methodology: MF, GT, DS, YY

Investigation: SR, SC, HJ, YY, HS, TG, BS, GT, MF, SR

Visualization: HS, MF, HJ, YY, GT

Supervision: MF, GT, DS

Writing—original draft: MF, HJ

Writing—review & editing: HJ, MF, GT, AP

## Competing interests

Dr. Davey Smith DMS has consulted for FluxErgy Inc, Kiadis Pharmaceuticals, Bayer Pharmaceuticals, Linear Therapies, Matrix Biomed, Model Medicines, VxBiosciences, and Brio Clinical. Dr. Marcelo Freire has consulted for Mars Wrigley, Bristle Health.

## Data and materials availability

All data are available in the main text or the supplementary materials. The raw proteomic data that support the findings of this study are shown in the source data file (ftp://MSV000086946@massive.ucsd.edu) at the Global Natural Products Social (GNPS) molecular networking depository via the Mass spectrometry Interactive Virtual Environment (MASSIVE).

## Notes

### Summary of Updates

Figure 1 was updated for better description of disease course; Abstract/introduction/discussion was updated to clarify our research aims One author was added; contributed revision of paper significantly

