## Supplementary material for "Persistent Immune and Clotting Dysfunction Detected in Saliva and Blood Plasma after COVID-19": Suppl tables and a figure

**This PDF file includes:**

Fig. S1
Tables S1 to S2

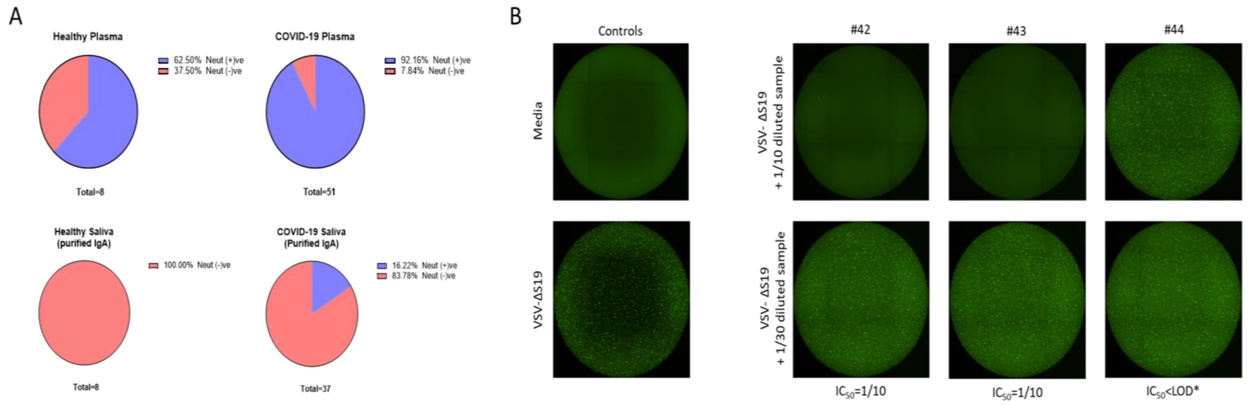


**Fig. S1.**

**Pseudo-virus neutralization by plasma and saliva samples**

Protection potency of plasma and saliva sample was measured by neutralizing activity to VSV pseudoviruses bearing S protein (rVSV-GFPΔG*Spike). Percentage of plasma or saliva samples with neutralizing activity was depicted as in pie chart (A). Representative images of inhibition of GFP expression by saliva samples were displayed in (B).

**Table S1.**

Demographic analysis of human participants.

| | | Healthy | COVID-19 |
|---|---|---|---|
| **DEMOGRAPHICS & SOCIO-ECONOMIC CULTURAL DATA** | | | |
| **No. of Subjects-n** (% of Total Subjects) | | 13 | 34 |
| **Gender** (% of total withing group) | Male | 7 (53.84%) | 13 (38.24%) |
| | Female | 6 (46.16%) | 21 (61.76%) |
| **Age (years)-mean ± SD** (min-max) | | 41.85 ± 5.79 (33- 55) | 45 ± 11.90 (20-72) |
| **DISEASE INDICES** | | | |
| **Classification of clinical symptoms** (% of total withing group) | health | 13 (100%) | - |
| | Asymptomatic | - | 2 (5.88%) |
| | Mild | - | 19 (55.88%) |
| | Moderate | - | 11 (32.35%) |
| | Severe | - | 2 (5.88%) |
| **Days from clinical onset-mean ± SD** (min-max) | | - | 51.28 ± 19.84 (20-90) |

**Table S2A.**

**Significant correlations among all immunoglobulin subtypes in healthy controls.**

| Immunoglobulins | Pearson's R | p-value | Immunoglobulins | Pe |
|---|---|---|---|---|
| RBD Saliva IgA vs.NP Saliva IgA | 0.897 | 0.003 | S1 Plasma IgA vs. S2 Plasma IgM | [illegible] |
| RBD Saliva IgA vs.NL63 Saliva IgM | 0.727 | 0.041 | S2 Plasma IgA vs. S1 Plasma IgM | [illegible] |
| S1 Saliva IgA vs.NP Saliva IgA | 0.880 | 0.004 | S2 Plasma IgA vs. NP Plasma IgM | [illegible] |
| S1 Saliva IgA vs.NL63 Saliva IgM | 0.863 | 0.006 | S2 Plasma IgA vs. NL63 Plasma IgM | [illegible] |
| S1 Saliva IgA vs. S1 Plasma IgA | -0.747 | 0.033 | NP Plasma IgA vs. S1 Plasma IgG | [illegible] |
| S2 Saliva IgA vs. S1 Plasma IgG | -0.798 | 0.018 | NL63 Plasma IgA vs.NP Plasma IgG | [illegible] |
| NP Saliva IgA vs.NL63 Saliva IgA | 0.897 | 0.003 | RBD Plasma IgG vs. NL63 Plasma IgG | [illegible] |
| NP Saliva IgA vs.NL63 Saliva IgM | 0.902 | 0.002 | RBD Plasma IgG vs. S1 Plasma IgM | [illegible] |
| NP Saliva IgA vs. S1 Plasma IgA | -0.768 | 0.026 | RBD Plasma IgG vs.NP Plasma IgM | [illegible] |
| NP Saliva IgA vs.NP Plasma IgA | -0.714 | 0.047 | RBD Plasma IgG vs. RBD Plasma IgM | [illegible] |
| NP Saliva IgA vs. S1 Plasma IgG | -0.868 | 0.005 | RBD Plasma IgG vs. S2 Plasma IgM | [illegible] |
| NL63 Saliva IgA vs.NL63 Saliva IgM | 0.727 | 0.041 | NL63 Plasma IgG vs.NP Plasma IgM | [illegible] |
| S1 Saliva IgG vs. RBD Plasma IgA | 0.977 | 0.004 | NL63 Plasma IgG vs. RBD Plasma IgM | [illegible] |
| NP Saliva IgG vs. S1 Plasma IgM | 0.771 | 0.025 | NL63 Plasma IgG vs. S2 Plasma IgM | [illegible] |
| NP Saliva IgG vs. S2 Plasma IgM | 0.645 | 0.084 | NL63 Plasma IgG vs.NL63 Plasma IgM | [illegible] |
| NP Saliva IgG vs.NP Plasma IgM | 0.771 | 0.025 | RBD Plasma IgM vs. NP Plasma IgM | [illegible] |
| NL63 Saliva IgM vs. S1 Plasma IgG | -0.199 | 0.001 | RBD Plasma IgM vs.NL63 Plasma IgM | [illegible] |
| NL63 Saliva IgM vs. S1 Plasma IgA | -0.379 | 0.005 | S1 Plasma IgM vs. S2 Plasma IgM | [illegible] |
| NL63 Saliva IgM vs.NP Plasma IgA | -0.200 | 0.043 | S1 Plasma IgM vs. NL63 Plasma IgM | [illegible] |
| S1 Plasma IgA vs.NP Plasma IgA | 0.222 | 0.015 | S2 Plasma IgM vs. NP Plasma IgM | [illegible] |
| S1 Plasma IgA vs. S1 Plasma IgG | -0.237 | 0.017 | S2 Plasma IgM vs.NL63 Plasma IgM | [illegible] |
| S1 Plasma IgA vs. RBD Plasma IgM | 0.118 | 0.036 | | |

**Table S2B.**

**Significant correlations among all immunoglobulin subtypes in convalescent COVID-19.**

| Immunoglobulins | Pearson's R | p-value | Immunoglobulins | Pearson's R | p-value |
|---|---|---|---|---|---|
| RBD Saliva IgA vs. RBD Plasma IgA | 0.701 | <0.0001 | NP Saliva IgG vs. RBD Plasma IgG | 0.701 | <0.0001 |
| RBD Saliva IgA vs. RBD Saliva IgM | 0.558 | <0.0001 | NP Saliva IgG vs. NL63 Plasma IgG | 0.701 | <0.0001 |
| RBD Saliva IgA vs. S1 Saliva IgM | 0.515 | <0.0001 | NP Saliva IgG vs. S2 Plasma IgG | -0.359 | 0.020 |
| RBD Saliva IgA vs. NL63 Saliva IgG | 0.502 | 0.001 | NP Saliva IgG vs. S1 Plasma IgG | -0.356 | 0.021 |
| RBD Saliva IgA vs. S1 Plasma IgG | 0.488 | 0.001 | NL63 Saliva IgG vs. S1 Saliva IgM | 0.995 | 0.000 |
| RBD Saliva IgA vs. RBD Plasma IgM | 0.314 | 0.043 | NL63 Saliva IgG vs. RBD Plasma IgA | 0.776 | <0.0001 |
| RBD Saliva IgA vs. S2 Plasma IgM | 0.314 | 0.043 | NL63 Saliva IgG vs. RBD Plasma IgM | 0.646 | <0.0001 |
| S1 Saliva IgA vs. NL63 Plasma IgM | 0.506 | 0.001 | NL63 Saliva IgG vs. S2 Plasma IgM | 0.646 | <0.0001 |
| S1 Saliva IgA vs. RBD Plasma IgM | 0.504 | 0.001 | NL63 Saliva IgG vs. RBD Saliva IgM | 0.606 | <0.0001 |
| S1 Saliva IgA vs. S2 Plasma IgM | 0.504 | 0.001 | NL63 Saliva IgG vs. NL63 Saliva IgM | -0.512 | <0.0001 |
| S1 Saliva IgA vs. S1 Plasma IgM | 0.433 | 0.004 | NL63 Saliva IgG vs. S2 Saliva IgM | -0.425 | 0.004 |
| S1 Saliva IgA vs. NL63 Saliva IgA | 0.319 | 0.037 | NL63 Saliva IgG vs. NP Plasma IgA | -0.333 | 0.031 |
| S1 Saliva IgA vs. NP Saliva IgG | 0.313 | 0.044 | RBD Saliva IgM vs. S1 Saliva IgM | 0.552 | 0.000 |
| S1 Saliva IgA vs. S1 Plasma IgA | 0.309 | 0.046 | RBD Saliva IgM vs. RBD Plasma IgM | 0.422 | 0.005 |
| S2 Saliva IgA vs. NP Saliva IgA | 0.294 | 0.000 | RBD Saliva IgM vs. S2 Plasma IgM | 0.422 | 0.005 |
| S2 Saliva IgA vs. NL63 Saliva IgM | 0.010 | 0.001 | RBD Saliva IgM vs. S1 Plasma IgG | 0.385 | 0.012 |
| S2 Saliva IgA vs. NP Plasma IgA | 0.030 | 0.003 | S1 Saliva IgM vs. RBD Plasma IgA | 0.776 | <0.0001 |
| S2 Saliva IgA vs. S2 Saliva IgM | -0.025 | 0.010 | S1 Saliva IgM vs. RBD Plasma IgM | 0.646 | <0.0001 |
| NP Saliva IgA vs. S2 Saliva IgM | 0.390 | 0.000 | S1 Saliva IgM vs. S2 Plasma IgM | 0.646 | <0.0001 |
| NP Saliva IgA vs. NL63 Saliva IgA | -0.028 | 0.001 | S1 Saliva IgM vs. NL63 Saliva IgM | -0.528 | <0.0001 |
| NP Saliva IgA vs. NP Saliva IgG | -0.023 | 0.001 | S1 Saliva IgM vs. S2 Saliva IgM | -0.500 | 0.001 |
| NP Saliva IgA vs. S2 Saliva IgG | 0.195 | 0.002 | S1 Saliva IgM vs. NP Plasma IgA | -0.333 | 0.031 |
| NP Saliva IgA vs. RBD Saliva IgG | 0.030 | 0.015 | S2 Saliva IgM vs. NL63 Saliva IgM | 0.559 | <0.0001 |
| NP Saliva IgA vs. NP Saliva IgM | 0.245 | 0.027 | S2 Saliva IgM vs. NP Plasma IgA | 0.563 | <0.0001 |
| NP Saliva IgA vs. RBD Plasma IgA | -0.181 | 0.048 | S2 Saliva IgM vs. RBD Plasma IgA | -0.448 | 0.003 |
| NL63 Saliva IgA vs. NL63 Plasma IgA | 0.196 | <0.0001 | S2 Saliva IgM vs. NP Saliva IgM | 0.404 | 0.008 |
| NL63 Saliva IgA vs. RBD Plasma IgG | 0.196 | <0.0001 | S2 Saliva IgM vs. RBD Plasma IgM | -0.387 | 0.011 |
| NL63 Saliva IgA vs. NL63 Plasma IgG | 0.196 | <0.0001 | S2 Saliva IgM vs. S2 Plasma IgM | -0.387 | 0.011 |

| | | | | | |
|---|---|---|---|---|---|
| NL63 Saliva IgA vs. RBD Saliva IgG | 0.372 | <0.0001 | S2 Saliva IgM vs. S2 Plasma IgG | 0.320 | 0.039 |
| NL63 Saliva IgA vs. S2 Saliva IgG | 0.469 | 0.001 | NP Saliva IgM vs. NL63 Saliva IgM | 0.365 | 0.017 |
| NL63 Saliva IgA vs. S2 Plasma IgG | -0.124 | 0.020 | NP Saliva IgM vs. S2 Plasma IgA | -0.311 | 0.045 |
| NL63 Saliva IgA vs. S1 Plasma IgG | -0.099 | 0.021 | NL63 Saliva IgM vs. NP Plasma IgA | 0.912 | <0.0001 |
| RBD Saliva IgG vs. S1 Saliva IgG | -0.300 | 0.002 | NL63 Saliva IgM vs. S2 Plasma IgG | 0.471 | 0.002 |
| RBD Saliva IgG vs. S2 Saliva IgG | 0.475 | 0.002 | NL63 Saliva IgM vs. RBD Plasma IgA | -0.411 | 0.007 |
| RBD Saliva IgG vs. NP Plasma IgG | 0.220 | 0.004 | NL63 Saliva IgM vs. RBD Plasma IgM | -0.389 | 0.011 |
| RBD Saliva IgG vs. NP Saliva IgM | 0.051 | 0.023 | NL63 Saliva IgM vs. S2 Plasma IgM | -0.389 | 0.011 |
| RBD Saliva IgG vs. NL63 Plasma IgA | 0.701 | 0.037 | NL63 Saliva IgM vs. S1 Plasma IgA | 0.360 | 0.019 |
| RBD Saliva IgG vs. RBD Plasma IgG | 0.701 | 0.037 | NP Plasma IgA vs. S2 Plasma IgG | 0.011 | 0.004 |
| RBD Saliva IgG vs. NL63 Plasma IgG | 0.701 | 0.037 | RBD Plasma IgA vs. RBD Plasma IgM | 0.504 | 0.001 |
| RBD Saliva IgG vs. S1 Plasma IgM | -0.155 | 0.050 | RBD Plasma IgA vs. S2 Plasma IgM | 0.504 | 0.001 |
| S1 Saliva IgG vs. NP Plasma IgA | 0.449 | 0.003 | RBD Plasma IgA vs. S1 Plasma IgG | 0.344 | 0.026 |
| S1 Saliva IgG vs. NL63 Saliva IgM | 0.392 | 0.009 | S1 Plasma IgA vs. NL63 Plasma IgA | -0.042 | 0.004 |
| S1 Saliva IgG vs. S1 Plasma IgM | 0.396 | 0.009 | S1 Plasma IgA vs. RBD Plasma IgG | -0.042 | 0.004 |
| S1 Saliva IgG vs. S2 Plasma IgG | 0.374 | 0.015 | S1 Plasma IgA vs. NL63 Plasma IgG | -0.042 | 0.004 |
| S1 Saliva IgG vs. NL63 Plasma IgM | 0.365 | 0.017 | S1 Plasma IgA vs. NP Plasma IgA | -0.280 | 0.005 |
| S1 Saliva IgG vs. NP Saliva IgG | -0.322 | 0.037 | NP Plasma IgA vs. S2 Plasma IgG | 0.439 | 0.004 |
| S1 Saliva IgG vs. NP Plasma IgM | 0.299 | 0.055 | NL63 Plasma IgA vs. NP Plasma IgG | 0.361 | 0.019 |
| S2 Saliva IgG vs. NP Plasma IgG | 0.547 | <0.0001 | S1 Plasma IgG vs. NP Plasma IgG | 0.315 | 0.042 |
| S2 Saliva IgG vs. NP Saliva IgG | 0.495 | 0.001 | NP Plasma IgG vs. NL63 Plasma IgG | 0.361 | 0.019 |
| S2 Saliva IgG vs. NL63 Plasma IgA | 0.364 | 0.018 | RBD Plasma IgM vs. NL63 Plasma IgM | 0.622 | <0.0001 |
| S2 Saliva IgG vs. RBD Plasma IgG | 0.364 | 0.018 | RBD Plasma IgM vs. S1 Plasma IgM | 0.589 | <0.0001 |
| S2 Saliva IgG vs. NL63 Plasma IgG | 0.364 | 0.018 | S1 Plasma IgM vs. NL63 Plasma IgM | 0.720 | <0.0001 |
| NP Saliva IgG vs. NL63 Plasma IgA | 0.701 | <0.0001 | S1 Plasma IgM vs. NP Plasma IgM | 0.481 | 0.001 |
| NP Saliva IgG vs. RBD Plasma IgG | 0.701 | <0.0001 | S2 Plasma IgM vs. NL63 Plasma IgM | 0.622 | <0.0001 |
| NP Saliva IgG vs. NL63 Plasma IgG | 0.701 | <0.0001 | | | |
